# functionInk: An efficient method to detect functional groups in multidimensional networks reveals the hidden structure of ecological communities

**DOI:** 10.1101/656504

**Authors:** Alberto Pascual-García, Thomas Bell

**Affiliations:** Department of Life Sciences. Silwood Park Campus. Imperial College London, Ascot, United Kingdom; Institute of Integrative Biology. ETH-Zürich, Zürich, Switzerland

**Keywords:** Multiplex networks, community detection, modules, guilds, mutualistic networks, trophic networks, microbial networks

## Abstract

1. Complex networks have been useful to link experimental data with mechanistic models, and have become widely used across many scientific disciplines. Recently, the increasing amount and complexity of data, particularly in biology, has prompted the development of multidimensional networks, where dimensions reflect the multiple qualitative properties of nodes, links, or both. As a consequence, traditional quantities computed in single dimensional networks should be adapted to incorporate this new information. A particularly important problem is the detection of communities, namely sets of nodes sharing certain properties, which reduces the complexity of the networks, hence facilitating its interpretation.
2. In this work, we propose an operative definition of “function” for the nodes in multidimensional networks, and we exploit this definition to show that it is possible to detect two types of communities: i) modules, which are communities more densely connected within their members than with nodes belonging to other communities, and ii) guilds, which are sets of nodes connected with the same neighbours, even if they are not connected themselves. We provide two quantities to optimally detect both types of communities, whose relative values reflect their importance in the network.
3. The flexibility of the method allowed us to analyze different ecological examples encompassing mutualistic, trophic and microbial networks. We showed that by considering both metrics we were able to obtain deeper ecological insights about how these different ecological communities were structured. The method mapped pools of species with properties that were known in advance, such as plants and pollinators. Other types of communities found, when contrasted with external data, turned out to be ecologically meaningful, allowing us to identify species with important functional roles or the influence of environmental variables. Furthermore, we found the method was sensitive to community-level topological properties like the nestedness.
4. In ecology there is often a need to identify groupings including trophic levels, guilds, functional groups, or ecotypes.The method is therefore important in providing an objective means of distinguishing modules and guilds. The method we developed, *functionInk* (functional linkage), is computationally efficient at handling large multidimensional networks since it does not require optimization procedures or tests of robustness. The method is available at: HTTPS://GITHUB.COM/APASCUALGARCIA/FUNCTIONINK.

## 1 Introduction

Networks have played an important role in the development of ideas in ecology, particularly in understanding food webs [1], and flows of energy and matter in ecosystems [2]. However, modern ecological datasets are becoming increasingly complex, notably within microbial ecology, where multiple types of information (taxonomy, behaviour, metabolic capacity, traits) on thousands of taxa can be gathered. A single network might therefore need to integrate different sources of information, leading to connections between nodes representing relationships of different types, and hence with different meanings. Advances in network theory have attempted to develop tools to analyse these more sophisticated networks, encompassing ideas such as multiplex, multilayer, multivariate networks, reviewed in [3]. There could therefore be much value in extending complex networks tools to ecology in order to embrace these new concepts.

Broadly speaking, a network represents how a large set of entities *share* or *transmit* information. This definition is intentionally empty-of-content to illustrate the challenges we face in network analysis. For instance, a network in which information is shared may be built relating genes connected if their sequence similarity is higher than a certain threshold. In that case, we may capture how their similarity diverged after an evolutionary event such as a gene duplication. On the other hand, networks may describe how information is transmitted, as in an ecosystem in which we represent how biomass flows through the trophic levels or how behavioural signals are transmitted among individuals. We aim to illustrate with these examples that, when building networks that consider links of different nature (e.g. shared vs. transmitted information) or different physical units (e.g. biomass vs. bits), it is not straightforward to extrapolate methods from single-dimensional to multidimensional complex networks.

In this paper we aim to address a particularly relevant problem in complex networks theory, namely the detection of “communities”[4] when the network contains different types of links. In addition, we are interested in finding a method with the flexibility to identify different types of communities. This is motivated by the fact that, in ecology, it is recognized that communities may have different topologies [5] and often an intrinsic multilayer structure [6]. We aim to detect two main types of communities. Firstly, the most widely adopted definition of community is the one considering sets of nodes more densely connected within the community than with respect to other communities, often called *modules* [7]. An example in which modules are expected is in networks representing significant co-occurrences or segregations between microbial species, when these relationships are driven by environmental conditions. Since large sets of species may simultaneously change their abundances in response to certain environmental variables [8, 9], this results in large groups of all-against-all co-occurring species, and between-groups segregations. Secondly, we are interested in finding nodes sharing a similar connectivity pattern even if they are not connected themselves. An example comes from networks connecting consumers and their resources, when communities are determined looking for consumers sharing similar resources preferences. This idea is aligned with the classic Eltonian definition of niche, which emphasizes the functions of a species rather than their habitat [10]. We call this second class of communities *guilds*, inspired by the ecological meaning in which species may share similar ways of exploiting resources (i.e. similar links) without necessarily sharing the same niche (not being connected themselves), emphasizing the functional role of the species [11]. Consequently, guilds may be quite different to modules, in which members of the same module are tightly connected by definition. The situation in which guilds are prevalent is known as disassortative mixing [12], and its detection has received comparatively less attention than the “assortative” situation (which results in modules) perhaps with the exception of bipartite networks [13, 14].

There are many different approximations for community detection in networks, summarized in [15]. However, despite numerous advances in recent years, it is difficult to find a method that can efficiently find both modules and guilds in multidimensional networks, and that is able to identify which is the more relevant type of community in the network of interest. This might be because there is no algorithm that can perform optimally for any network [16], and because each type of approximation may be suited for some networks or to address some problems but not for others, as we illustrate below.

Traditional strategies to detect modules explore trade-offs in quantities like the betweeness and the clustering coefficient [7], as in the celebrated Newman-Girvan algorithm [17]. Generalizing the determination of modules to multidimensional networks is challenging. Consider, for instance, that a node A is linked with a node B and this is, in turn, linked with a node C, and both links are of a certain type. If A is then linked with node C with a different type of link, should the triangle ABC be considered in the computation of the clustering coefficient? One solution proposed comes from the consideration of stochastic Laplacian dynamics running in the network [18], where the permanence of the informational fluxes in certain regions of the network reflects the existence of communities. This approximation has been extended to consider multilayer networks [19], even if there are modules defined in different layers that highly overlap, hence defining communities (combination of modules) across layers [20]. A fundamental caveat for these methods is that the links must have a clear interpretation for how their presence affects informational fluxes. Returning to the above example, if the links AB and BC represent mutualistic interactions and the link AC represents a competitive one, can a random walk follow the link AC when this interaction does not describe a flux of biomass between the species but rather a disruption in the flux of biomass of AB and BC?

A related approach searches for modules using an optimization function that looks for a partitioning in a multilayer network that maximizes the difference between the observed model and a null model which considers the absence of modules [13]. This strategy can be applied to multidimensional networks, but raises questions such as which is the appropriate null model, and how to determine the coupling between the different layers defining the different modes of interaction [21]. In addition, since these approximations focus on the detection of modules, they neglect the existence of guilds or other network structures.

Regarding the search of guilds, this problem has received notable attention in social sciences following the notion of structural equivalence. Two nodes are said to be structurally equivalent if they have the same connectivity in the network [22]. The connectivity may be defined either analyzing if two nodes share the same neighbours, if two nodes are connected with neighbors of the same type even if they are not necessarily the same (following some preassigned roles for the nodes, e.g. prey are structurally equivalent because are connected to predators), or a combination of both. Social agents often have an assigned role, which is why structural equivalence is particularly important in social networks.

An approximation that has exploited the idea of structural equivalence is stochastic blockmodelling [23], which considers generative models with parameters fitted to the observed network. The approach brings greater flexibility because different models can accommodate different types of communities [24]. Therefore, this approximation could be used to search for both modules and guilds [25]. There are, however, also important caveats to the approach, since it is an important challenge to determine whether the underlying assumptions of a particular block model is appropriate for the data being used [26]. Moreover, even when the model brings an analytically closed form, the estimation of the parameters may be computationally intractable [27], hence requiring costly optimality procedures or tests for robustness [28].

In this work, we build on the idea of structural equivalence noting that a node belonging to either a guild or a module is, in both cases, structurally equivalent to the other nodes in its community. This observation was acknowledged in social sciences in the definition of λ−communities [29], which are types of communities encompassing both modules and guilds, whose relevance has also been previously recognized in the ecological literature [5, 30]. From this observation, we wondered whether it is possible to find a similarity measure between nodes that quantifies their structural equivalence, even when different types of links are considered. We could then join nodes according to this similarity measure while monitoring whether the communities that are formed are guilds or modules. A similar approach was investigated by Yodzis *et al.* to measure trophic ecological similarity [31], but they did not identify an appropriate threshold for determining community membership (which they call “trophospecies”).

We have developed an approach that builds on these results and develops a method to determine objective thresholds for identifying modules and guilds in ecological networks. We show that a modification of the community detection method developed by Ahn et al. [32], leads to the identification of two quantities we call internal and external partition densities. For a set of nodes joined within a community by means of their structural equivalence similarity, the partition densities quantify whether their similarities come from connections linking them with nodes outside the community (external density) or within the community (internal density). Notably, our method generates maximum values for the two partition densities along the clustering, allowing us to objectively determine thresholds for the similarity measure in which the communities correspond to the definition of modules (for the internal density) and guilds (external density). Since the elements within both types of communities are structurally equivalent, modules and guilds can be understood as different kinds of *functional groups* –in the Eltonian sense– and this is the name we adopt here. We reserve the term “community” for a more generic use, because other types of communities beyond functional groups may exist, such as core-periphery structures [33].

We call our method functionInk (functional linkage), emphasizing how the number and types of links of a node determine its functional role in the network. We illustrate its use by considering complex ecological examples, for which we believe the notion of functional role is particularly relevant. We show in the examples that, by combining the external and internal partition densities, we are able to identify the underlying dominant structures of the network (either towards modules or towards guilds). Moreover, selecting the most appropriate community definition in each situation provides results that are comparable to state-of-the-art methods. This versatility in a single algorithm, together with its low computational cost to handle large networks, makes our method suitable for any type of complex, multidimensional network.

## 2 Methods

### Structural equivalence similarity in multidimensional networks

Our method starts by considering a similarity measure between all pairs of nodes that quantifies the fraction of neighbours connected with links of the same type that they share (Fig. 1). This is a natural definition of structural equivalence for multidimensional networks, which is agnostic to the specific information that the interaction carries. For simplicity, we present a derivation for a network that contains two types of links. We use undirected positive (+) interactions (e.g. a mutualism) and negative (−) interactions (e.g. competition) to illustrate the method, but these could be replaced by any two link types. Extending the method to an arbitrary number of link types is presented in the Suppl. Material. We call {*i*} the set of *N* nodes and {*e*_*ij*_} the set of *M* links in a network. We call *n*(*i*) the set of neighbours of *i*, that can be split into different subsets according to the types of links present in the network.

**Figure 1:**
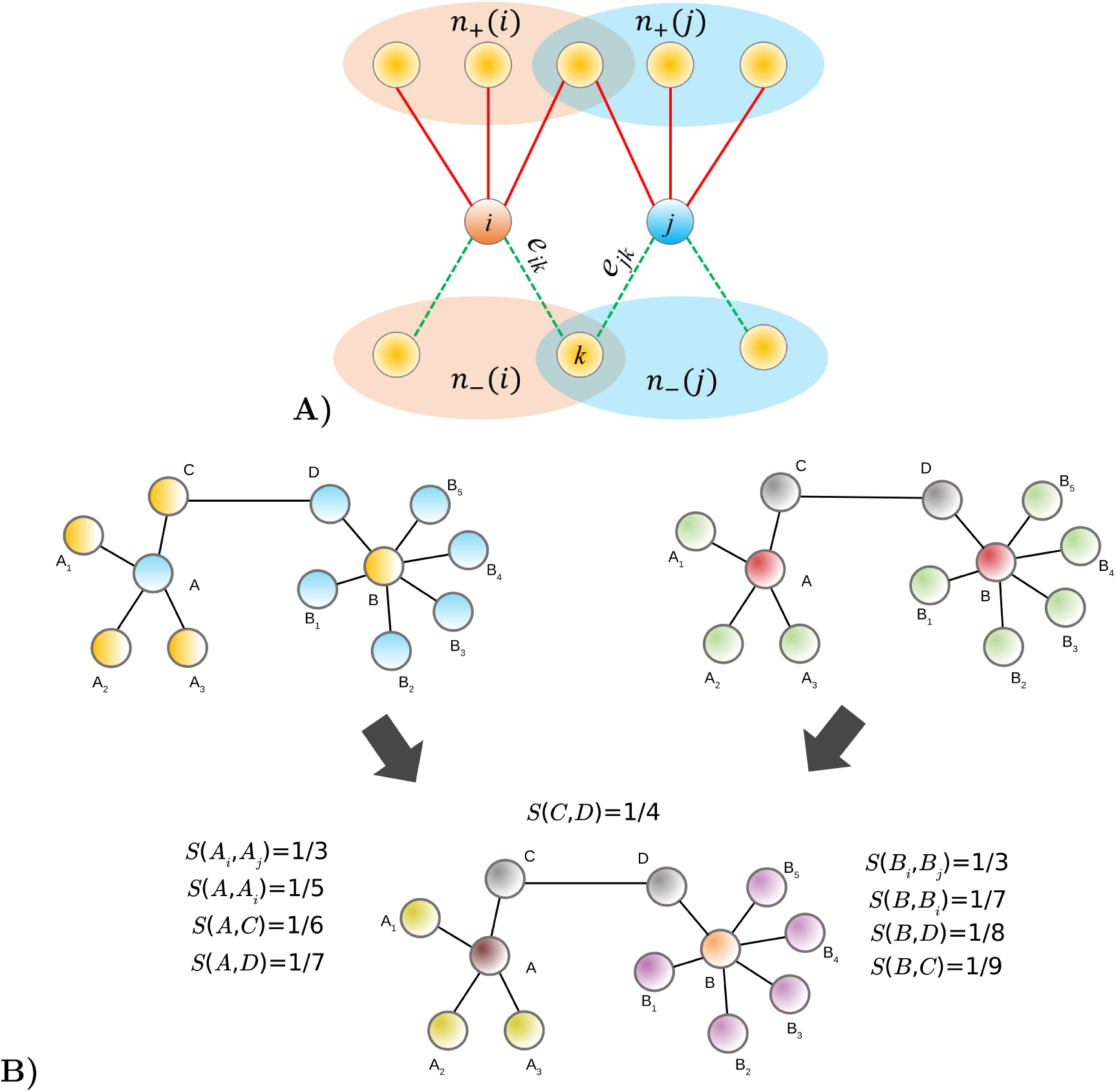
Illustration of the method. (A) The similarity between nodes *i* and *j* is computed considering the neighbours of each node and the types of interactions that link them. In this example, two types of link are shown: positive (+) interactions are solid links connecting the sets of neighbours *n*_+_(*i*) and *n*_+_(*j*). Negative (−) links are shown as dotted links connecting the sets of neighbours *n*_−_(*i*) and *n*_−_(*j*). Following Eq. 2, |*n*(*i*) ∩ *n*(*j*) | = 2 and |*n*(*i*) ∪ *n*(*j*)| = 8, which yields *S*^(J)^(*i, j*) = 2*/*8. If, for instance, *e*_*ik*_ changes from being - to +, the node *k* would no longer belong to the set *n*(*i*) ∩ *n*(*j*), being the new similarity: *S*^(J)^(*i, j*) = 1*/*8. In Ref. [32] the similarity computed in this way is assigned to the links *e*_*ik*_ and *e*_*jk*_. (B) Structural equivalence can be defined in different ways. In the top-left network we considered that blue and yellow colors encode *a priori* information describing the roles of the nodes. Identifying sets of nodes connected similarly to nodes with equivalent roles (i.e. the emphasis is on the roles and not on the specific neighbours, a situation called regular equivalence [34]) leads to two communities (the yellow and blue sets of nodes themselves), because every blue node is connected to a yellow one. The method of Guimerá and Amaral [33], determines communities focusing on their topological role (top-right network) by identifying central (A and B), peripheral (A1-A3 and B1-B5) and connector nodes (C and D). functionInk (bottom network) defines communities by joining nodes with approximately the same neighbours and, if there are roles for the nodes, these can be incorporated defining link types (one type for each pair of roles connected, in the example only one type is needed). All non-zero Jaccard similarities of the example are shown. Clustering these similarities will lead to different partitions and, stopping at *S*^(J)^ = 1*/*4, communities being the intersection of those found in the above networks are obtained, highlighting the potential to identify communities considering both the roles and topological features. Figure adapted from [33].

For two types of links, we split the set of neighbours linked with the node *i* into those linked through positive relationships, *n*_+_(*i*), or through negative relationships, *n*_−_(*i*); we follow a notation similar to the one presented in [32], but note that *n*(*i*) there denotes neighbours irrespective of the type of links. Distinguishing link types induces a division in the set of neighbours of a given node into subsets sharing the same link type, shown in Fig. 2A. More specifically, in the absence of link types we define the Jaccard similarity between two nodes *i* and *j* as:

**Figure 2:**
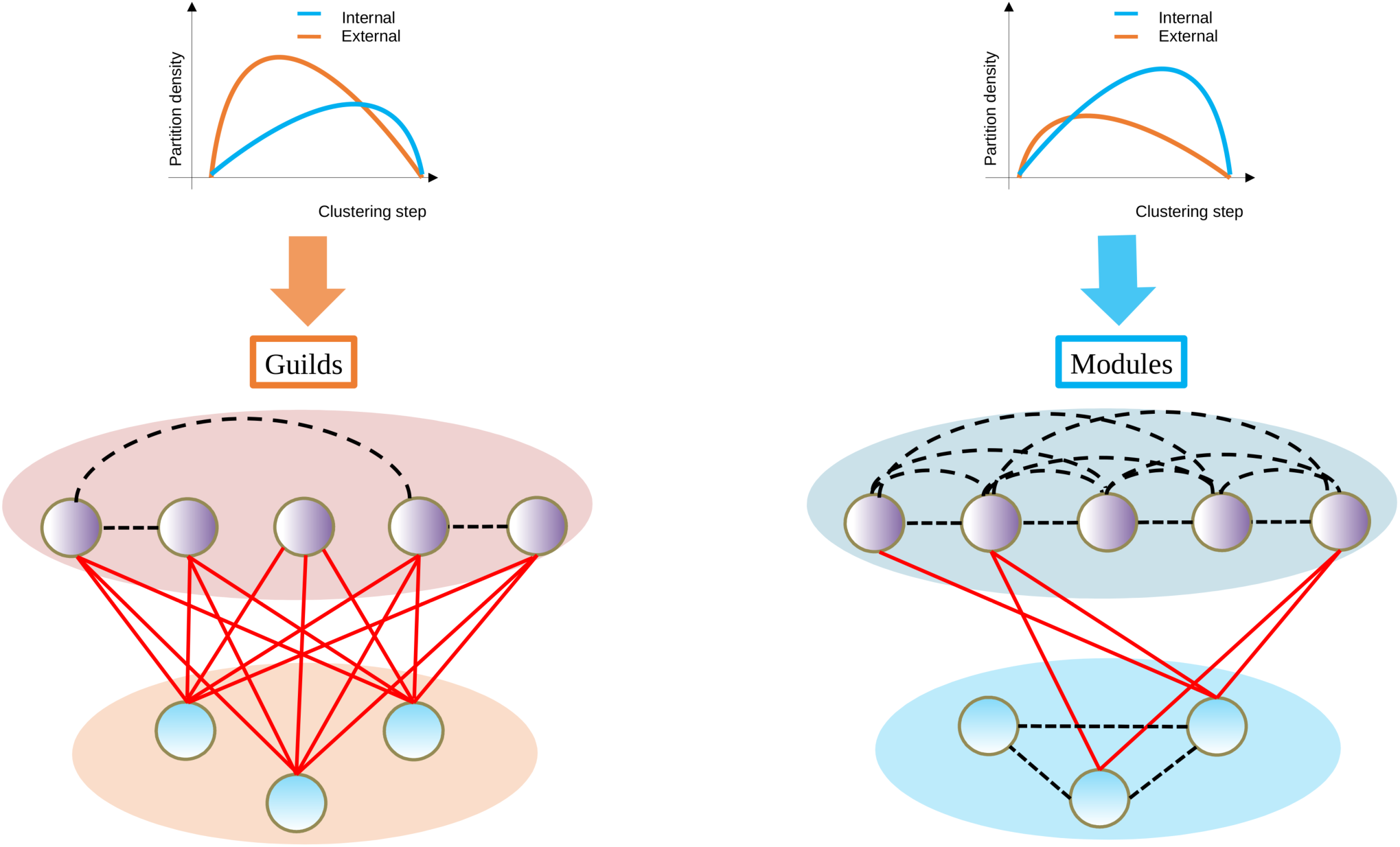
Definition of guilds and modules. For each set of nodes 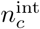 belonging to the same community *c* (nodes within the same shaded area) we consider the number of links within the community (black dashed links, called 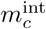 in the main text) out of the total number of possible internal links, to compute the internal partition density (see upper curves). We also computed the external partition density, which is the density of links connecting nodes external to the community (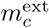, solid red lines linking nodes belonging to different communities) out of the total number of possible external links. We call guilds the communities determined at the maximum of the external partition density, and modules those found at the maximum of the internal partition density. The relative value of the external and internal partition densities allow us to estimate which kind of community dominates the network. In the example, guilds dominate the network on the left, and modules dominates the network on the right.

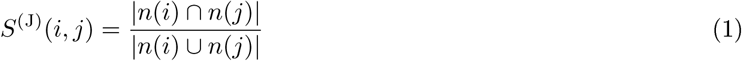

where |·| is the cardinality of the set (the number of elements it contains). This metric was shown to lead to clusters of species that are more consistent with cophenetic clustering than other alternatives [31], and generalizing this expression to multiple attributes is achieved simply by differentiating the type of neighbours depending on the types of connections. For two attributes (see Suppl. Material for an arbitrary number of attributes) this leads to

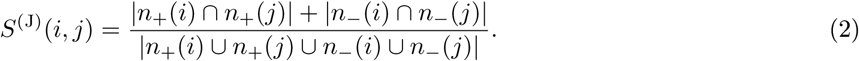

Accounting for the weight of the links can be made with the generalization of the Jaccard index provided by the Tanimoto coefficient [35], *S*^(T)^(*i, j*), presented in Suppl. Material.

Finally, we introduce a modification to the above definition of *S*^(J)^ to account for the particular case in which *i* and *j* are only connected between themselves, i.e. they do not share any neighbours according to the above definition. This is problematic because we want to distinguish this situation from the one in which they do not share any nodes, for which we get *S*(*i, j*) = 0. We resolve this situation by considering that a node is its own neighbour, in which case two nodes only connected between themselves would yield *S*(*i, j*) = 1. However, we note that this would also be the value between two nodes that are connnected and that also share all neighbours (a motif known as a clique), irrespective of the number of neighbours they share, because the similarity measure saturates. We argue that this situation is unsatisfactory because there is stronger evidence that two nodes are structurally equivalent when they share connections and creating transitive motifs, since transitivity is a key property in the definition of equivalence classes [36]. The situation can be resolved by using the convention that, for two connected nodes, the intersection set is reduced by one, i.e. |*n*(*i*) ∩ *n*(*j*)| → |*n*(*i*) ∩ *n*(*j*)| − 1. This convention has the interesting property that, for cliques, increasing the number of nodes involved also increases the similarity between its members, resulting in an upper bound of one and a lower bound of 1/2 (a 2-node clique). In addition, two connected nodes that share neighbors but are not connected themselves have a smaller difference in the similarity compared to nodes within cliques that share the same number of neighbors, thus facilitating the identification of guilds. In Fig. 1 we illustrate the computation of this similarity with a simple example.

### Identification of communities through clustering and similarity cut-offs

Once the similarity between nodes is computed, the next objective is to define and identify structurally equivalent communities. As explained in the Introduction, there are different possible definitions of structural equivalence, illustrated in Fig. 1B. In the figure is shown how the similarity metric proposed together with an agglomerative clustering to join nodes in communities, encapsulates these different notions of structural equivalence. A critical question, however, is how to objectively determine the threshold to stop the clustering ([37])?

This question is often addressed by iteratively “partitioning” the network into the distinct communities, and monitoring each partition with a function having a well defined maximum or minimum that determines the threshold of the optimal partition. In [32], the authors proposed to join links of a network according to a similarity measure between the links with an agglomerative clustering, and to monitor the clustering with a quantity called the partition density. The partition density is the weighted average across communities of the number of links within a community out of the total possible number of links (which depends on the number of nodes in the community). We re-considered the method of Ahn et al. [32] (which was originally defined over partitions of links, see Suppl. Material), to work over partitions of nodes, and we developed two partition densities, with two distinct meanings. To develop these measures we noted that, when joining nodes into a cluster, we are concluding that these nodes share (approximately) the same neighbours connected with the same type of links, but the nodes joined may or may not be connected between them. We therefore redefined the partition density so that it distinguishes between the contribution to the link density arising from the connections *within* a community from connections shared with with external nodes *between* communities.

Formally, given a node *i*, we differentiate neighbours that are within the same community (*n*^int^(*i*), where int stands for “interior”) from neighbours that are in different communities (*n*^ext^(*i*)), hence *n*(*i*) = *n*^int^(*i*) ∪ *n*^ext^(*i*) (Fig. 2). For a singleton (a community of size one) *n*^int^(*i*) = {*i*} and *n*^ext^(*i*) = ∅. Similarly, the set of links *m*(*i*) connecting the node *i* with other nodes can also be split into two sets: the set connecting the node with neighbours within its community *m*^int^(*i*), and those connecting it with external nodes *m*^ext^(*i*). This distinction was also considered in other context (called the problem of coloring nodes [38]).

Therefore, for each partition of nodes into *T* communities our method identifies, for each community *c*, the total number of nodes it contains, 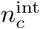, and the total number of links connecting these nodes 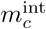. In addition, it computes the total number of nodes in other communities that have connections to the nodes in the community, 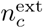, through a number of links 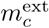. Clearly, to identify 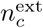 neighbours, at least 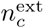 links are required and thus an increasing number of links in excess, 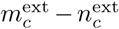 are necessary to obtain an increasing contribution to the similarity of the nodes in the community through external links (however, this is not a sufficient condition, see Suppl. Material). In this way, a relevant quantity to characterize a community is the fraction of links in excess out of the total possible number 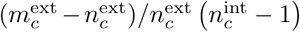. We note this calculation does not take into account multiple link types. The weighted average of this quantity through all communities leads to the definition of external partition density:

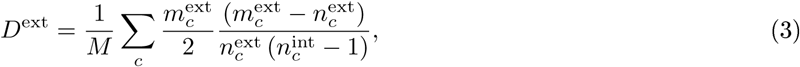

where *M* is the total number of links. We now follow a similar reasoning to consider a necessary condition to obtain an increasing contribution to the similarity of the nodes through the internal links (see Suppl. Material). We acknowledge that in a community created by joining nodes through the similarity measure we propose, it may happen that 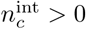 even if 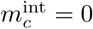. Therefore, any link is considered a link in excess, leading to the following expression for the internal partition density, which quantifies the fraction of internal links in excess out of the total:

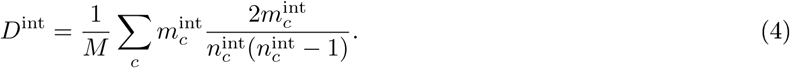

Finally, we define the total partition density as the sum of both internal an external partition densities:

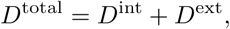

and hence, if all the fractions in *D*^int^ and *D*^ext^ are equal to one, i.e. all possible links in excess are realized, *D*^total^ equals to one. Since at the beginning of the clustering the communities have a low number of members, most of the contribution towards *D*^total^ comes from *D*^ext^ while, in the last steps, where the communities become large, *D*^int^ will dominate. All three quantities will reach a maximum value along the clustering (for the internal it could be at the last step) and, if one of them clearly achieves a higher value, it will be indicative that one type of functional group is dominant in the network. If that is the case, the maximum of *D*^total^ –which is always larger or equal to max(max(*D*^int^), max(*D*^ext^))−, will be at a clustering step close to the step in which the dominating quantity peaks. If neither *D*^ext^ nor *D*^int^ clearly dominates, *D*^total^ will peak at an intermediate step between the two partial partition densities maxima, suggesting that this intermediate step is the best candidate of the optimal partition for the network. Communities determined at this intermediate point where they can be both guilds and modules will be called, generically, functional groups.

## 3 Results

### Plant-pollinator networks

To illustrate the use of the method we start analyzing a synthetic example. In ecological systems, species are often classified into communities according to their ecological interactions, such as in mutualistic networks of flowering plants and their animal pollinators. These networks are characterized by intra and interspecific competition within both the pool of plants and the pool of animals, and by mutualistic relationships between plants and animals, leading to a bipartite network.

To investigate the performance of our method and, in particular, the influence of the topological properties into the partition density measures, we generated a set of artificial mutualistic networks with diverse topological properties, following the method presented in [39]. For the mutualistic interactions, we focused on two properties: the connectance *κ*_mut_, which is the fraction of observed interactions out of the total number of possible interactions, and the nestedness *ν* as defined in [40] (see Methods), which codifies the fraction of interactions that are shared between two species of the same pool, averaged over all pairs of species. We selected these measures for their importance in the stability-complexity debate in mutualistic systems [39], and the similarity between the nestedness (which, in the definition we adopt here, represents the mean ecological overlap between species) and the notion of structural equivalence we considered. For the competition matrices, we considered random matrices with different connectances, *κ*_comp_, since it is difficult to estimate direct pairwise competitive interactions experimentally, and they are frequently modeled with a mean field competition matrix.

We verified that in all networks the set of plants and animals are joined in the very last step of the clustering irrespective of the clustering method used, a result that must follow construction. As expected, the curves monitoring the external and internal partition densities depends on the properties of the networks. We illustrate this finding in Fig. 3, where we have selected two networks with contrasting topological properties. One of the networks has high connectance within the pools and low connectance and nestedness between the pools. The internal partition density peaks at the last step minus one (i.e. where the two pools are perfectly separated) consistent with the definition of modules, where the intra-modules link density is higher than the inter-modules link density. On the other hand, the second network has intra-pool connectance equals to zero, and very high connectance and nestedness between the pools (see Fig. 3). We selected a *κ*_comp_ = 0 for simplicity in the network representation, but similar results are obtained for low values of *κ*_comp_, see for instance Suppl. Fig. 10. In this second network (see Fig. 3, right panel), only the external partition density peaks and, at the maximum, the communities that we identified clearly reflect the structural equivalence of the nodes members in terms of their connectance with nodes external to the group, as we expect for the definition of guilds. The ecological information retrieved for guilds is clearly distinct from the information retrieved for the modules, the former being related to the topology of the network connecting plants and animals. We observe that guilds identify specialist species clustered together, which are then linked to generalists species of the other pool: a structure typical of networks with high nestedness.

**Figure 3:**
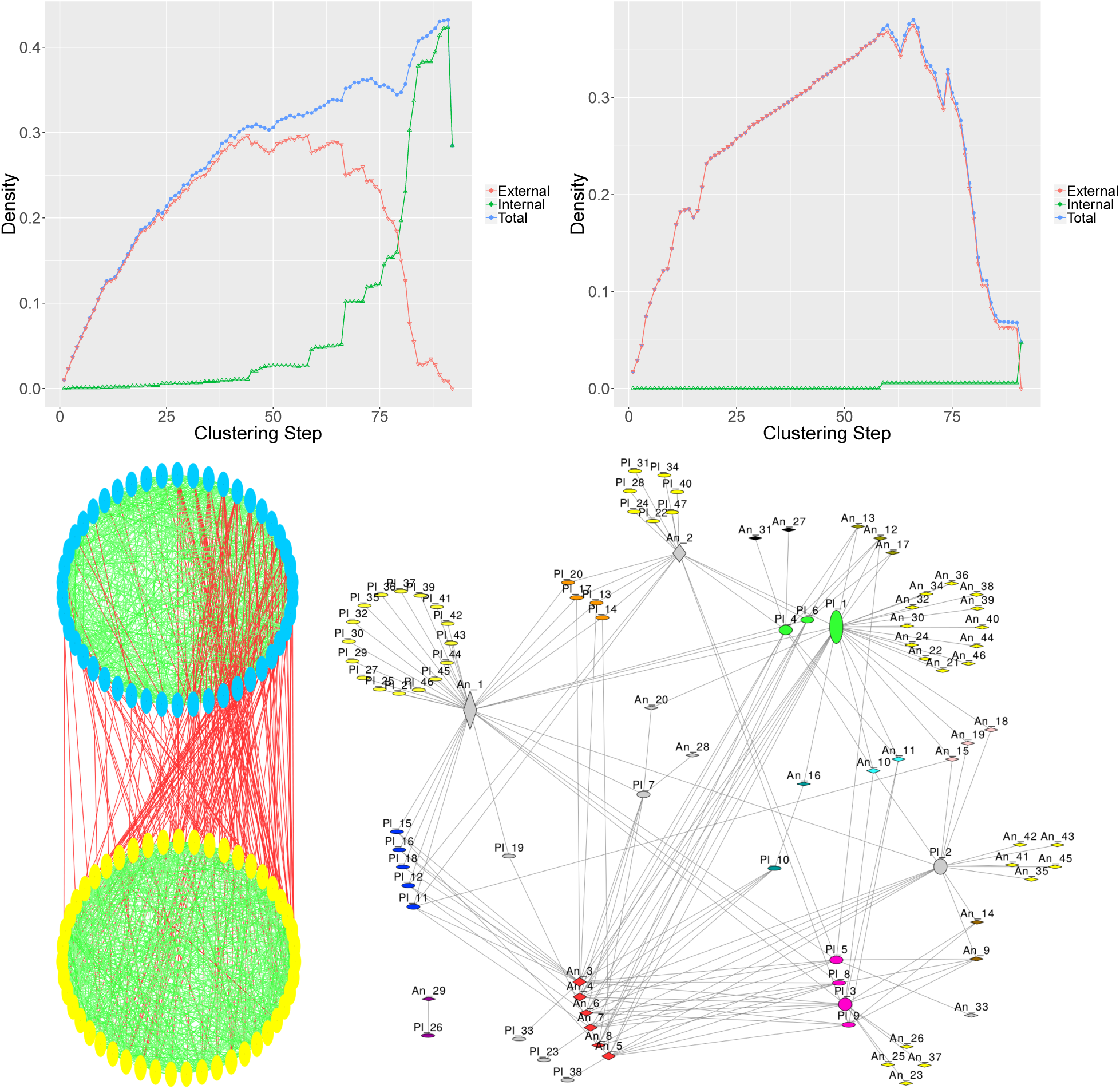
Analysis of synthetic mutualistic networks. (Top left) Partition densities for a network with *κ*_comp_ = 0.5, nestedness *ν* = 0.15 and *κ*_mut_ = 0.08 and (top right) for a network with *κ*_comp_ = 0, nestedness *ν* = 0.6 and *κ*_mut_ = 0.08. The high density of competitive links in the first network makes the internal partition density dominate, leading to two modules representing the plant-pollinator pools (bottom left network), while reducing the density of competitive links to zero in the second network makes the external partition density to dominate, finding guilds (bottom right, with plants labeled “Pl” and animals labeled “An”). The small increase in the internal partition density for this network at step 59 is due to two specialist species joined at that step (animal 29 and plant 56, shown at the bottom left of the network). Nodes are colored according to their functional group in both networks although, in the network finding guilds (bottom right), specialist species are yellow, single species communities are gray, and the size of the nodes is proportional to their betweeness.

The method identified several interesting guilds and connections between them. For instance, generalists Plant 1, Animal 1 and Animal 2 (and to a lesser extent Plant 2) have a low connectivity between them but, being connected to many specialists, determine a region of high vulnerability, in the sense that a directed perturbation over these species would have consequences for many other species. This is confirmed by the high betweeness of these nodes (proportional to the size of the node in the network). In addition, the algorithm is able to identify more complex partitions of nodes into communities. As an example of this, Animal 16 (turquoise) is split from Animals 10 and 11 (cyan), which form a second community, and from Animals 15, 18 and 19 (light pink) that are joined into a third community, despite of the subtle connectivity differences between these six nodes. Finally, it also detects communities of three or more species that have complex connectivity patterns which, in this context, may be indicative of functionally redundant species (e.g. red and blue communities).

Examples with other intermediate properties are analyzed in the Suppl. Figs. 8 and 9. Broadly speaking, either the internal or the total partition density maximum peaks at the last step minus one, allowing for detection of the two pools of species. Nevertheless, the method fails to find these pools in situations in which the similarity between members of distinct pools is comparable to the similarity of members belonging to the same pool. This may be the case if the connectances are small (see Suppl. Fig. 10). The relative magnitude of the external vs. internal partition density depends on the connectance between the pools of plants and animals and on the connectance within the pools, respectively (see Suppl. Fig. 8). Interestingly, networks for which the nestedness is increased keeping the remaining properties the same generated an increase in the external partition density (see Suppl. Fig. 9). These examples illustrate how the external partition density is sensitive to complex topological properties, in particular to an increase in the dissasortativity of the network, as expected when guilds are dominant.

### Trophic networks

We tested our method in a comprehensive multidimensional ecological network of 106 species distributed in trophic layers with approximately 4500 interactions, comprising trophic and non-trophic interactions (approximately 1/3 of the interactions are trophic) [41]. This network was analyzed looking for communities extending a stochastic blockmodelling method [12] to deal with different types of interactions [41]. The estimation of the parameters of the model through an Expectation-Maximization algorithm requires controling the influence of random starting conditions since each initial condition may lead to a different result, and hence is needed to test the robustness of the results. Here we show that, in this example, our method is comparable with this approximation, and it has the advantage of being deterministic. Moreover, the simplicity of the method allows us to handle large networks with arbitrary number of types of links and to evaluate and interpret the results, as we show in the following.

Our method finds a maximum for the internal density when there are only three communities. Previous descriptions of the network identified three trophic levels in the network (Predators, Herbivores and Basal species). The latter are further subdivided into subgroups like (e.g. Kelps, Filter feeders), and there are some isolated groups like one Omnivore and Plankton. To match these subgroups we observed that the total partition density reaches a maximum close to the maximum of the external partition density (step 69) and maintains this value along a plateau until step 95 (see Suppl. Fig. 11). We analyzed results at both clustering thresholds finding that, at step 95, we obtain modules with a good agreement with the trophic levels, shown in Fig. 4. On the other hand, at step 69 we find a larger number of communities, some of which fit the definition of modules and others the definition of guilds (see Fig. 4).

**Figure 4:**
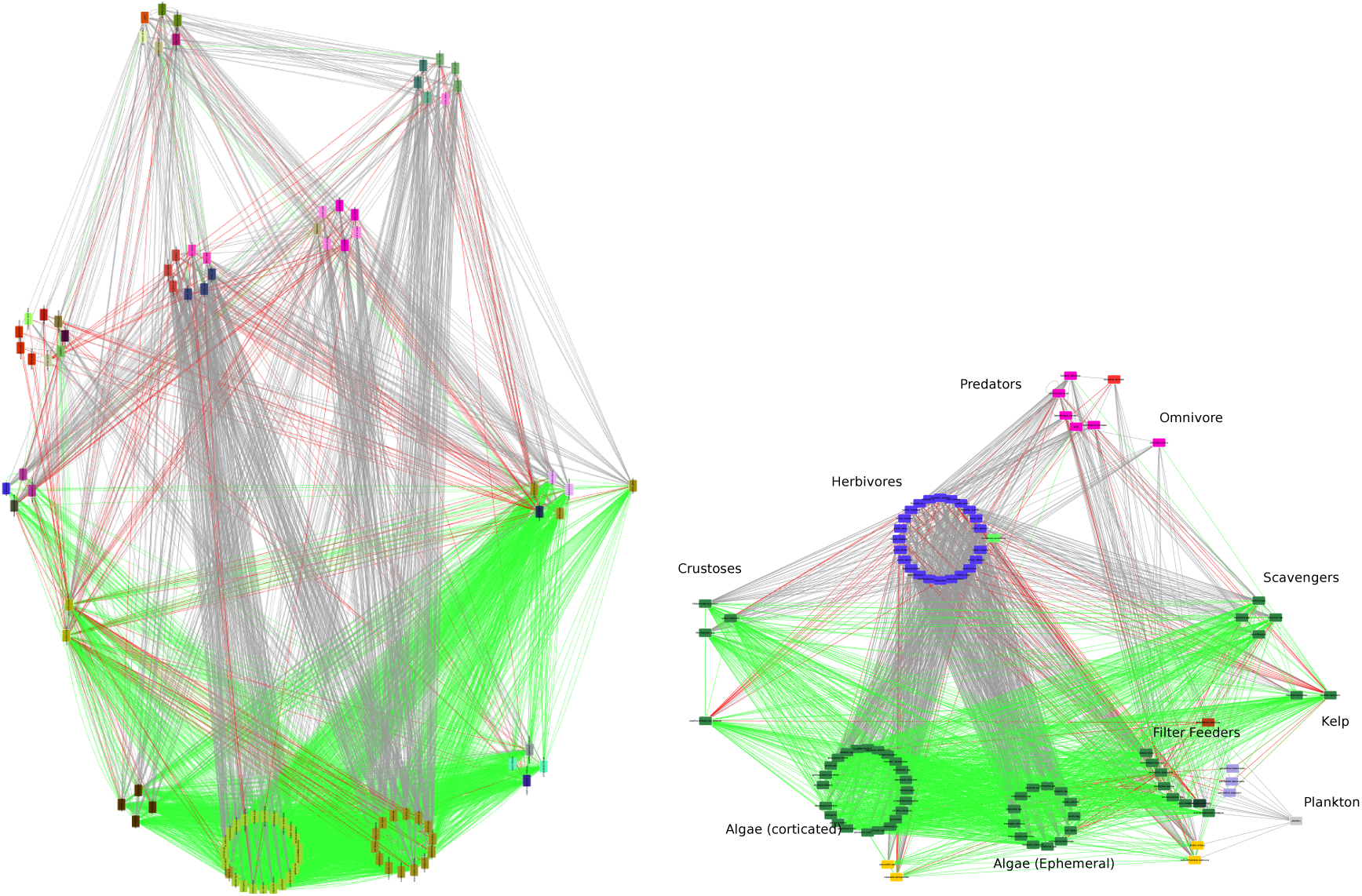
Determination of guilds and modules in a large trophic network. Trophic networks with links representing trophic (gray), non-trophic positive (red), and negative (green) interactions. (Left) Nodes are grouped according to the classification found in [41] (reference classification), and colored by the guilds found with functionInk at the maximum of the external partition density. (Right) Nodes are grouped according to the trophic levels and colored by the modules found by functionInk (see Main Text for details). The modules separate the three main trophic levels: predators, herbivores and basal species, further separating some of them into subgroups, such as filter feeders and plankton, which is an orphan module.

To shed some light on the information obtained from this second network, we compared the classification obtained by Kefi *et al.* [41] (in the following reference classification) and our method. We computed several similarity metrics comparing the classification we obtained at each step of the agglomerative clustering with functionInk and the reference classification (see Methods). In Fig. 5, we show that the similarity between both classifications is highly significant (Z-score > 2.5) and is maximized when the external partition density is also maximized, i.e. at step 69. This is particularly apparent for the Wallace 01, Wallace 10 and Rand indexes (see Fig. 6 and Suppl. Fig. 12). Notably, communities in the reference classification were also interpreted as functional groups in the same sense proposed here [41].

**Figure 5:**
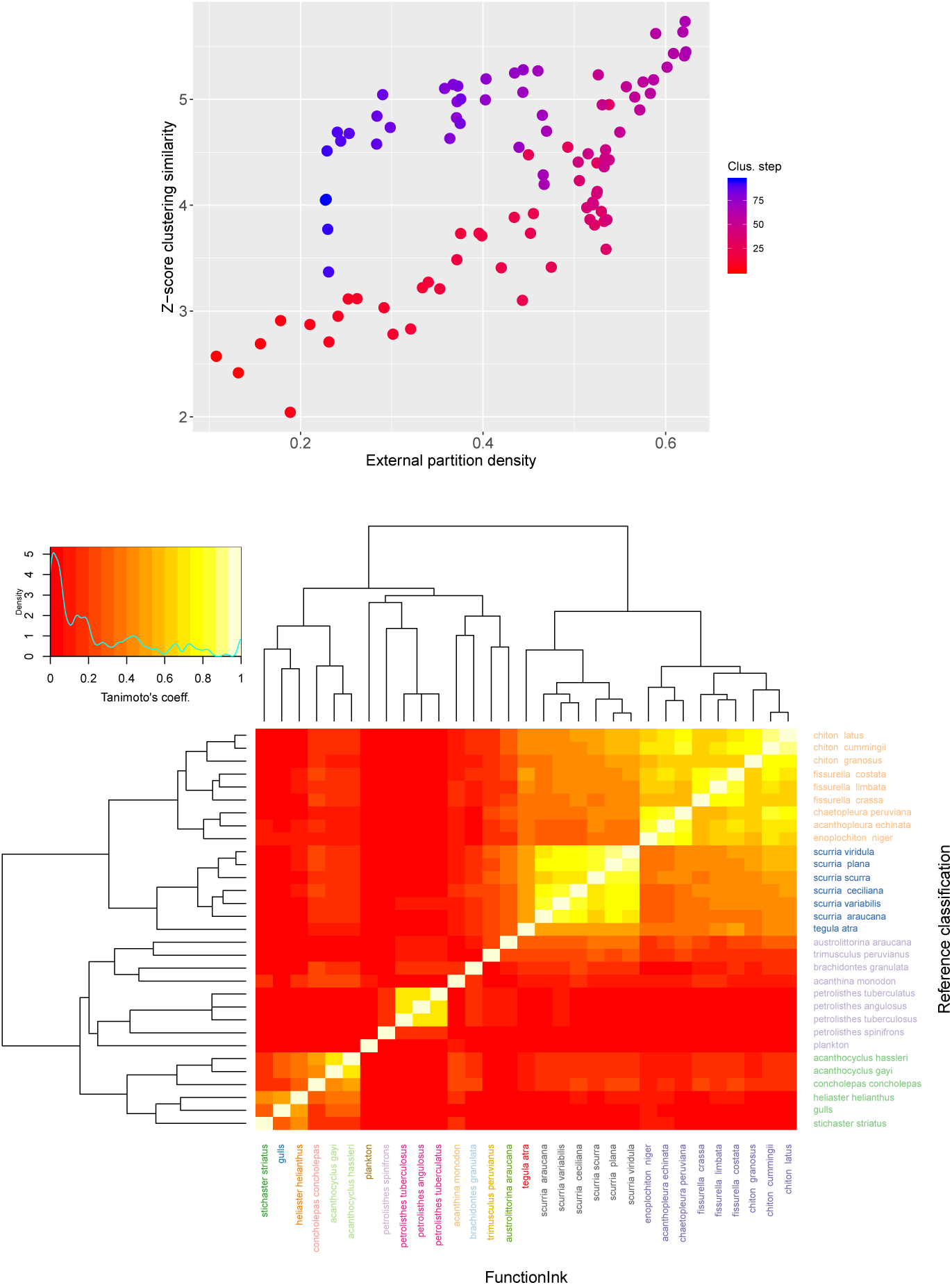
Comparison between the reference classification in the trophic network and functionInk. (Top) Z-score of the Wallace 10 index [42], measuring the similarity between the reference classification and the functionInk method at each clustering step. The similarity with the reference classification (see Main Text) is maximized around the maximum of the external partition density. (Bottom) Comparison of communities 1, 4, 7 and 9 in the reference classification, whose members were classified differently by functionInk. Colors in the names of species in rows (columns) represent community membership in the reference (functionInk) classifications. The heatmap represents the values of the Tanimoto coefficients, and the dendrograms are computed using Euclidean distance and clustered with complete linkage. Both classifications are generallyconsistent with the dendrograms, but with functionInk finding finer clusters.

**Figure 6:**
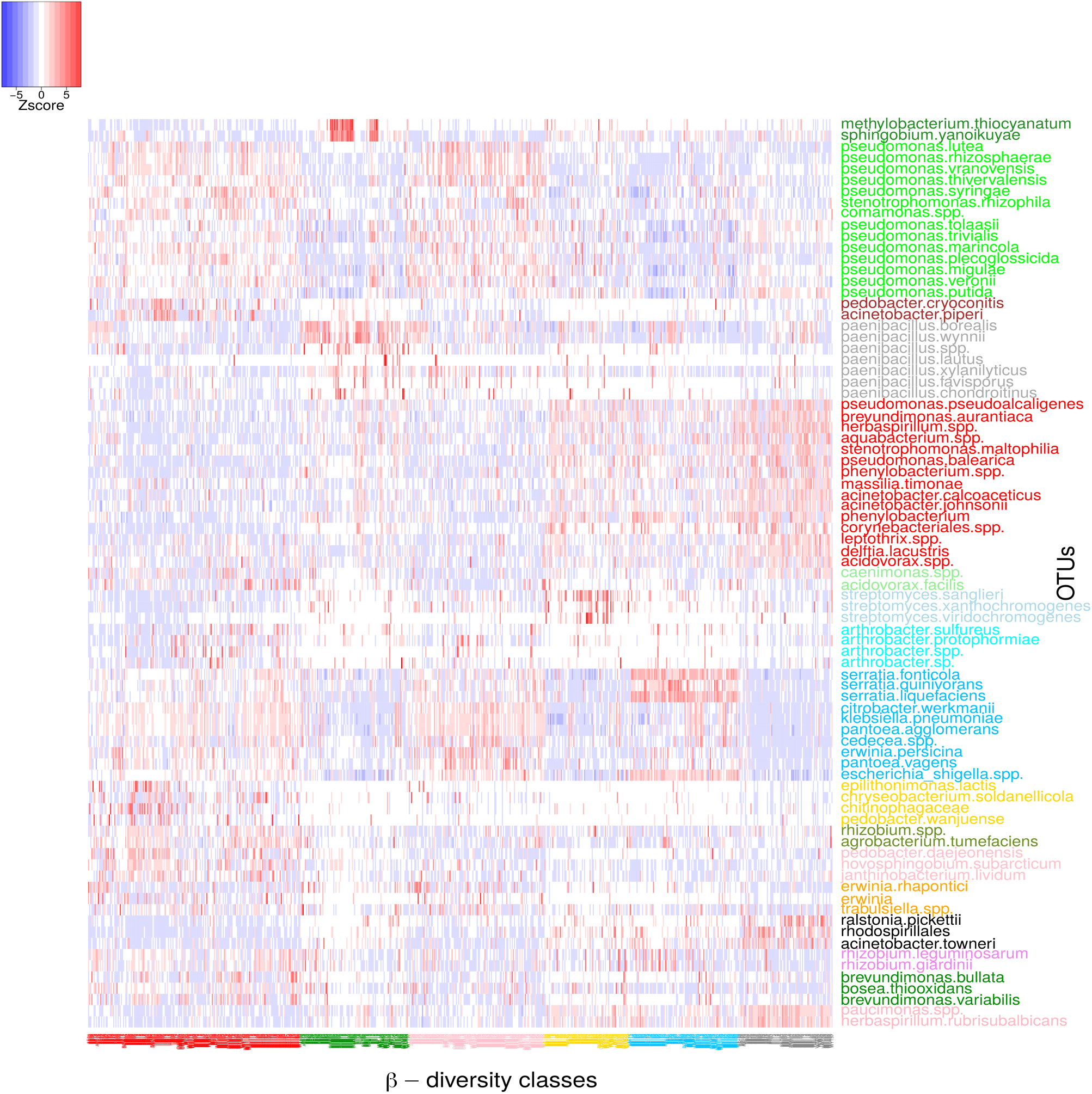
Comparison between *β*−diversity classes and functional groups in a microbial network. Heatmap representing the z-score of the log-transformed abundances of the OTUs (see Methods). Species are colored according to their functional group membership obtained at the maximum of the total partition density. Samples are colored according to one of the six community classes found in [43] after optimal clustering with a *β*−diversity distance. Orphan clusters were excluded except for 5 *Paenibacillus* species (characteristic of the green class) that were added to the functional group formed by *Paenibacillus* borealis and *Paenibacillus wynii*. The heatmap blocks show segregation and co-occurrence between modules, further mapping the *β*−diversity classes.

Nevertheless, there are some discrepancies between both classifications. In particular, although there is a complete correspondence between the two largest communities in both classifications, there are a number of intermediate communities in the reference classification whose members are classified differently in our method. To illustrate these discrepancies, we plotted a heatmap of the Tanimoto coefficients of members of four communities of intermediate sizes containing discrepancies, showing their membership in both the reference and the functionInk classification with different colors (see Fig. 5). The dendrograms cluster rows and columns computing the Euclidean distance between their values. Therefore, these dendrograms are very similar to the method encoded in functionInk, and the communities must be consistent, representing a powerful way to visually inspect results. Indeed, the dendrograms are in correspondence with both functionInk and reference communities, but we observe some discrepancies. For instance, the community found by the reference classification containing several *Petrolishtes* species, joins species that have low similarity regarding the number and type of interactions as measured by the Tanimoto coefficients, while functionInk joins together the three species with high similarity, leaving aside the remainder species. Therefore, despite the methodological differences between both methods, the different classifications produce similar outcomes, but result in different sized clusters (a different cut-off in the dendrograms, with functionInk finding finer clusters). The advantage of funtionInk is then apparent in the simplicity of the method, which permits validation through visual inspection of the consistency of the classification.

### Microbial networks

We discuss a last example of increasing importance in current ecological research, which is the inference of interactions among microbes sampled from natural environments. We considered a large matrix with more than 700 samples of 16S rRNA operative taxonomic units (OTUs) collected from rain pools (water-filled tree-holes) in the UK [43, 44] (see Suppl. Material). We analyzed *β*−diversity similarity of the samples contained in the matrix with the Jensen-Shannon divergence metric [45], further classifying the samples automatically, leading to 6 disjoint clusters we call *β*−diversity-classes (i.e. clusters of samples, see Methods). Next, we inferred a network of significant positive (co-occurrences) or negative (segregations) correlations between OTUs using SparCC [46] (see Methods), represented in Suppl. Fig. 13. Applying functionInk to the network of inferred correlations, we aimed to understand the consistency between the results of functionInk (modules and guilds) and the *β*−diversity-classes. The rationale is that, by symmetry, communities determined from significant co-occurrences and segregations between OTUs should reflect the similarity and dissimilarity between the samples, hence validating the method.

Contrasting with the trophic network analyzed in the previous example, the external partition density brings a poor reduction of the complexity of the network (peaking after only 22 clustering steps), and the internal partition density is higher, hence suggesting a more relevant role for modules (see Suppl. Fig. 14). Differences in the three stopping criteria are shown in Suppl. Fig. 13, where two large modules are apparent, with a large number of intracluster co-occurrences (continuous links) and interclusters segregations (dotted links). Note that this is quite different to what is found in macroscopic trophic networks, where pools of species (e.g. prey) have within module competitive (segregating) interactions, while between-modules interactions can be positive (for predators) or negative (for prey).

There is reasonable agreement between the functional groups found at the maximum of the total partition density and the *β*−diversity-classes, shown in Fig. 6. Moreover, the detection of networks complements the information that *β*−diversity-classes provides, since it is possible to individuate the key players of these classes (see Suppl. Material). Notably, it was shown in [43] that the *β*−diversity-classes might be related to a process of ecological succession driven by environmental variation, the functional groups are likely driven by environmental preferences rather than by ecological interactions, likely explaining the large number of positive co-occurrences. This speaks against a naive interpretation of correlation networks in microbial samples as ecological interactions unless environmental preferences are under control [9, 8].

## Discussion

We presented a novel method for the analysis of multidimensional networks, with nodes with an arbitrary number of link types. We implemented the method adopting the definition of structural equivalence, which underlies both the similarity measure definition and the rationale behind both the clustering and our definition of partition densities. We selected a set-theoretic similarity measure quantifying the number of nodes that are shared with the same type of interaction, which we believe is a natural definition of structural equivalence for multidimensional networks, and that has the advantage that it does not make assumptions on how the information flows in the network, typical of approximations based on Laplacian dynamics (see e.g. [18, 19]). This allow us to join nodes simply by their similarity, with no need for specific assumptions about the network structure. Moreover, this similarity can also be naturally linked to two measures of nodes’ partitioning that allowed us to propose a clear differentiation between modules (determined by the maximum of the internal partition density) and guilds (determined by the maximum of the external partition density).

Beyond these technical advantages, we illustrated the versatility of functionInk using several ecological examples. The relative value between the internal and external partition density immediately yields information on whether the network is dominated by modules, guilds, or intermediate structures. This allows for increasing flexibility in the analysis of the networks, and for a more nuanced interpretation of network structure and species’ roles in the ecosystem. For both mutualistic and trophic networks, the internal partition density correctly finds the trophic layers, justifying the success of the original method [32]. Our extension recovered the functional groups as determined by Kefi et al. [41] through the external partition density, and the visual inspection reflects a good consistency with the definition we proposed for functional groups in terms of structural equivalence. Moreover, in the mutualistic networks, we showed that the functional groups discovered in this way was sensitive to changes to high-order topological properties such as the nestedness.

The analysis of the microbial network was dominated by modules rather than guilds. Interestingly, these modules had intra-cluster positive correlations, contrary to what would be expected in a macroscopic trophic network, where competitive interactions would be dominant between members of the same trophic layer. We selected in this example for further exploration the functional communities found at the maximum of the total partition density, with some groups having properties closer to those of guilds and others closer to modules. The communities that we identified were in good agreement with the functional communities found using *β*−diversity similarity [43], supporting the consistency of the method. Interestingly, it was found in [43] that similar *β*−diversity-classes were driven by environmental conditions. Although co-occurring more often in the same environment may be indicative of a higher probability of interaction [47], the most economical hypothesis is that they co-occur because they share similar environmental preferences, and hence it cannot be disentangled the type of interaction (if any) unless the environmental variables are under control [9, 8].

To finish, we highlight some limitations of the method. Firstly, it may have problems if the communities are highly overlapping [20, 32]. In these cases, it would be convenient to inspect the partition at the three classifications given by the different partition densities, since it is likely that overlapping communities are split in an earlier classification and then joined at later steps of the clustering. Another possibility is to combine it with the approximation proposed in Ref. [32], that has both compatible and, at the same time, complementary results (see Suppl. Material). To continue with, our approximation does not consider yet the case in which there are multiedges in the network, although real networks are typically very sparse and the probability of finding multiedges is small [26]. Finally, although the method might not be able to achieve the generality of other approximations aiming to find any arbitrary structure in the network [12, 24, 28], such approximations require either heuristics to find a solution for the parameters −and hence a unique optimal solution is not guaranteed−, or a computationally costly sampling of the parameter space. Our method relies on a deterministic method whose results are easily inspected, and its computational cost for a network with *N* nodes scales as *N* ^2^ for the similarity metric computation plus the clustering, which is order *N*. The method is freely available in the address (HTTPS://GITHUB.COM/APASCUALGARCIA/FUNCTIONINK) and, importantly, although we developed it with ecological networks in mind, it can be applied to any kind of network.

## Acknowledgments

We thank Michael Schaub for critical comments on an earlier version of the manuscript, and to Sebastian Bonhoeffer for his support. We also thank to two anonymous referees whose comments help us to improve the manuscript. The research was funded by a European Research Council starting grant (311399-Redundancy) awarded to T.B. T.B. was also funded by a Royal Society University Research Fellowship. APG was also funded by the Simons Collaboration: Principles of Microbial Ecosystems (PriME), award number 542381.

## Authors’ Contributions

APG conceived the project and designed methodology; APG performed the analysis; TB contributed data; APG and TB analysed the data; APG led the writing of the manuscript. Both authors contributed critically to the drafts and gave final approval for publication.

## Supplementary Methods and Results

### Generalization of the Jaccard and Tanimoto coefficients to an arbitrary number of link types

Consider a network with a set {*i*} of *N* nodes and a set {*e*_*ij*_} of *M* links. These links are classified into Ω types labeled with the index *α* = (1, …, Ω). These types would typically account for differential qualitative responses of the nodes properties due to the interactions. For example, if we consider that the nodes are species and the property of interest is the species abundances, the effect of cooperative or competitive interactions on the abundances can be codified using two different types of links: positive and negative. If these relations are inferred through correlations between abundances, we could use a quantitative threshold (for instance a correlation equal to zero) to split the links into positive and negative correlations. In general, we may use a number of qualitative attributes or quantitative thresholds in the weights of the links to determine different types of links.

We call *n*(*i*) the set of neighbours of *i*, and we split these neighbours into (at most) Ω different subsets according to the types of links present in the network. The Jaccard coefficient defined in Eq. 2 can be extended, considering similarities between nodes, as:

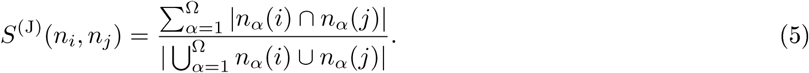

Accounting for the weight of the links can be made with the generalization of the Jaccard index provided by the Tanimoto coefficient [35]. We first introduce the method without differentiating between different types of neighbours. Consider the vector ***a***_*i*_ =(*Ã*_*i*1_, …, *Ã*_*ij*_) with

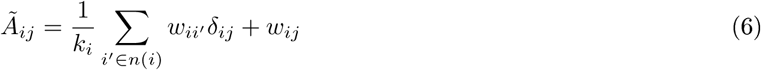

where *w*_*ij*_ is the weight of the link connecting the nodes *i* and *j, k*_*i*_ = |*n*(*i*)| and *δ*_*ij*_ is the Kronecker’s delta (*δ*_*ij*_ = 1 if *i* = *j* and zero otherwise). Determining the quantity *w*_*ij*_ = ***a***_*i*_***a***_*j*_ = ∑_*k*_ *Ã*_*ik*_*Ã*_*kj*_, the Tanimoto similarity is defined as

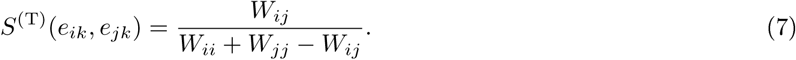

Working with link types requires a generalization of the above expression. Consider for the moment two types related with a positive *w*_*ij*_ > 0 or a negative *w*_*ij*_ < 0 weight of the links. The term *Ã*_*ii*_ = 1/*k*_*i*_∑*k*_*i*′_*w*_*ii*″_ is the average of the strengths of the links connected with node *i*, and it is desirable to keep this meaning when considering two types to properly normalize the Tanimoto similarity. This is simply achieved redefining *Ã*_*ij*_ as

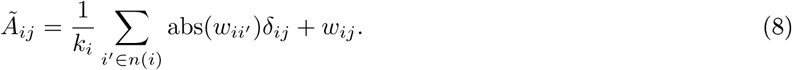

On the other hand, the similarity is essentially codified in the term *w*_*ij*_ that we now want to redefine to account for two types of interactions in such a way that only products *Ã*_*ik*_*Ã*_*kj*_ between terms with the same sign contribute to the similarity. This is achieved with the following definition, which generalizes the Tanimoto coefficient

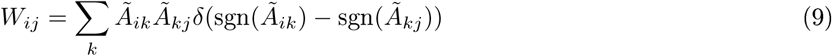

where sgn(·) is the sign function and *δ*(*a* − *b*) is the Dirac delta function (*δ*(*a* − *b*) = 0 if *a* ≠ *b*). Generalizing to an arbitrary number of types can be achieved by defining a variable *µ*_*ij*_ that returns the type of the link, i.e. *µ*_*ij*_ = *α* with *α* being a factor variable which, for the example of positive and negative links, is codified by the sign of the links’ weight. We generalize the expression in Eq. 9 as follows

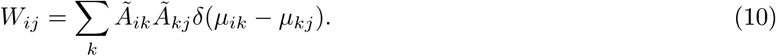

Finally, the generalization of the external and internal partition densities to consider multiple types of links simply requires us to correctly classify the neighbours of each node accounting for the different types 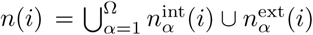. Similarly, the set of links *m*(*i*) connecting the node *i* with other nodes must be also split into sets according to the different types 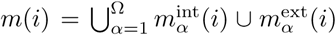. The expressions for the internal and external partition densities remain otherwise the same.

### Additional notes on the relation between the Jaccard similarity and the partition densities

In this section we aim to provide a more explicit relation between the Jaccard similarity and the partition density and, in particular, the notion of links in excess. We will show that the existence of links in excess is a neccessary condition to compute the similarity between the nodes in a cluster, and increases if the number of links in excess results in a higher similarity. Since the similarity can also be partitioned between external and internal components, this condition can be independently applied for the external and internal links in excess. We will finally discuss why these conditions are not sufficient.

Consider that the network has a single type of link (and we will make some precision below for the general case in which there are different types of links). In the computation of the partition densities, we inspect each community and we average across communities. Therefore, let us start considering a generic community *c*, and note that the neighbours of a node *i* are partitioned into those belonging to the same community, *n*^int^(*i*), and those belonging to other communities, *n*^ext^(*i*). The Jaccard similarity of two nodes within the same community can be expressed as

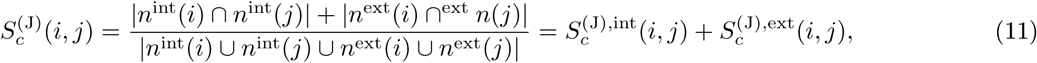

where we used a subindex *c* to stress the fact that *i, j* ∈ *c*. Hence, the mean similarity of the nodes in the community can also be split in two components

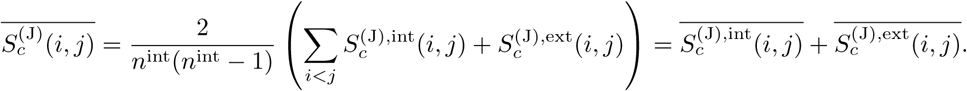

The sums 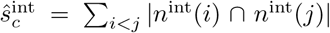 and 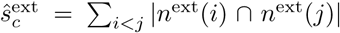 encode the contributions to the mean similarity of the community given by external and internal nodes, respectively. We aim to relate these terms to the links in excess used to compute the partition densities.

To establish this relationship, we note that the degree of any node in the network, |*n*(*i*)|, can also be partitioned between the links connecting to other nodes in the community it belongs to, |*n*^int^(*i*)|, and the links connecting to nodes from other communities, |*n*^ext^(*i*)|. Again, we will add a subindex, e.g. |*n*_*c*_(*i*)|, if there is any possible ambiguity about the identity of the community to which *i* belongs. In addition, we would like to specifically identify how the degree |*n*^ext^(*i*)| is partitioned with respect to each of the communities the node *i* is connected to, i.e. 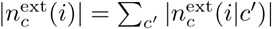, where 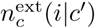 stands for the set neighbours of *i* ∈ *c* belonging to the communities *c*′ ≠ *c*. Similarly, we can now look at the degrees of the neighbours with respect to the community *c*, and we write 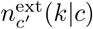 for the set of neighbours of *k* ∈ *c*′ belonging to the community *c*. Therefore, we can write the total

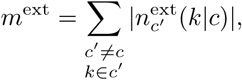

where the summatory runs for every element *k* ∈ *c*′ and every community *c*′ ≠ *c*.

In addition, it is evident that a node *k* in an external community *c*′ will contribute to the mean similarity of the nodes in community *c* if and only if its degree with respect to that community is at least 2, i.e. 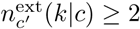, and thus it is immediate to relate 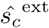 and 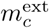:

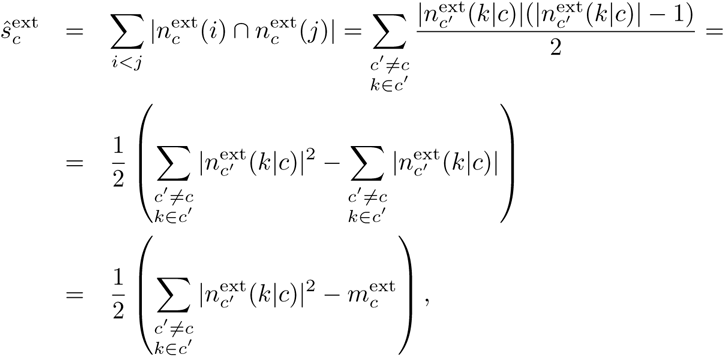

and therefore, if 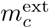 increases 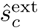 will increase (note that the argument of 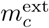 is the same that the quadratic term, but the latter will dominate the sum). We should still relate 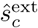 with the links in excess, which we defined as 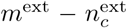 because we require at least one link with each neighbor to identify it as such. Therefore, we substract from each node one link, reducing its degree in one, and we again compute all the possible remaining pathways, that we substract from *ŝ*_*c*_:

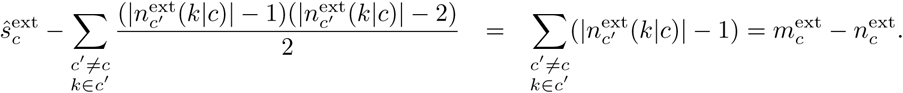

Note that the substracted term has the same arguments that we used for the computation of 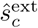 and that it must be smaller than 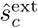. A similar reasoning can be followed for the internal partition density, although now we should note that all links contribute to the similarity because we followed a convention in which |*n*^int^(*i*) ∩ *n*^int^(*j*)| = 1 if *i* and *j* are linked, what leads to

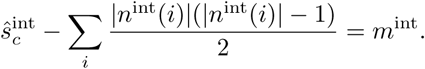

The main reason why an increase in the number of links in excess is not a sufficient condition for the similarity to increase is that, if the links in excess are of different types, they may not contribute to the similarity. This is a weakness of our approximation that would require reinspecting the types of connections between neighbours during the clustering procedure, and developing a procedure to solve ambiguities if the types are different. This is not required in the current implementation, in which the method is separated in two distinct problems, namely the computation of the similarities (where there is inspection of the neighbours and the types of connections) and the clustering, hence improving the computational efficiency. Since the clustering is performed using the *N* highest similarities (out of *N* (*N* − 1)*/*2 possible similarities, being *N* the number of nodes in the network), our choice is designed in a way in which nodes joined in a community are mostly connected to the same neighbours through the same type of links (otherwise the similarities should be low and the nodes will not be joined), which we confirmed in the examples shown is the article text. Hence, a more accurate partition density will likely not change the qualitative behaviour of the current functions, which have the advantage of being computationally efficient.

In addition, the relative contribution of 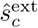 and 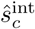 may not be strictly translated into the same relative contribution to the similarities because we are omitting in the above computation the normalization terms. For instance, it may happen that two nodes *i* and *j* are joined in a cluster and that, when an external node *k* joins the cluster, we have *ŝ*^ext^(*i, k*) > *ŝ*^ext^(*j, k*) and *S*^int^(*i, k*) > *S*^int^(*j, k*) (implying |*n*(*i*)| ≪ |*n*(*j*)|). This means that the neighbours shared by *i* and *k* and those shared between *j* and *k* substantially differ. However, since *k* has few neighbours and *i* and *j* were joined first, the inequality *S*(*i, j*) > max(*S*(*i, k*), *S*(*j, k*)) would be unlikely to be fulfilled (*n*(*i*) is large and |*n*(*i*) ∩ *n*(*j*)| is small since a fraction of the few neighbors that *j* has are shared with *k*). Therefore, again the hierarchical clustering helps avoid “pathological” situations, hence its importance in our approximation.

### Original definition of partition density and relation with functionInk

For completeness, we present the definition of partition density presented in [32] and a comparison with the new method. In short, Ahn *et al*. method starts building a similarity measure between any pair of links sharing one node in common. Two links will be similar if the nodes that these two links do not share have, in turn, similar relationships with any other node, shown in Fig. 1. From this similarity measure, links are clustered and an optimal cut-off for the clustering is found monitoring a measure called *partition density* (which in this paper relates to the *internal partition density*). The optimal classification found at the cut-off, determines groups of links that are similar because they connect nodes that are themselves similar in terms of their connectivity. Therefore, the nodes are classified indirectly, according to the groups that their respective links belong, and a node may not belong to a single community but to several communities if its links belong to different clusters. This is claimed to be an advantage with respect to other methods (in particular for high density networks) as membership to a single cluster is not enforced. At every step of the clustering it is obtained a partition *P* = *P*_1_, …, *P*_*C*_ of the links into *C* subsets. For every subset, the number of links is *m*_*c*_ = |*P*_*c*_| and the number of nodes that these links are connecting is 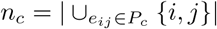. The density of links for the cluster *C* is then

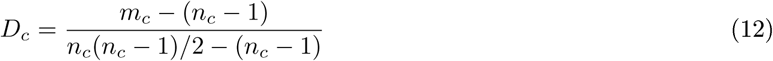

where the normalization considers the minimum (*n*_*c*_ − 1) and maximum (*n*_*c*_(*n*_*c*_ − 1)*/*2) number of links that can be found in the partition. The difference with respect to Eq. 4, is that a term 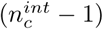 is now subtracted. The reason is that, in Ahn *et al*. method, clustering with links implies that two nodes in the same cluster must share links. But, according to our definition of function, two nodes may be structurally equivalent even if there is no interaction between them.

The partition density *D* is then given by the average of the density of links for all the partitions, weighted by the number of links

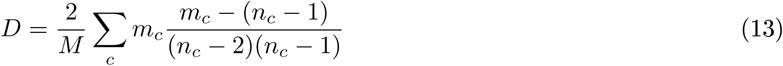

where *M* is the total number of links. It was shown that when using agglomerative clustering, this function achieves a maximum which determines the optimal partition [32].

functionInk shifts the attention from links back to nodes (see Fig. 1A). There are a number of reasons justifying this shift:

- Working with nodes is more natural, thus favoring the interpretation of the communities. From a biological perspective, we are interested in the function of nodes. For instance, the definition of “niche” is straightforward for nodes, justifying both the similarity measure definition and the rational behind both the clustering and the definition of partition densities we proposed. In particular, focusing on nodes allowed us to identify both modules and guilds.
- Inspecting and interpreting results is more convenient working with node partitions. For instance, most of the network visualization programs can easily separate communities defined over nodes partitions but not over links partitions.
- Monitoring changes in communities defined over node partitions becomes easier (even more if there are changes in the network such as addition/removal of nodes or links), because the number of links scales as *N* ^2^ while the number of nodes scales *N* (being *N* the number of nodes), so it may be expected a more profound change in the links partitioning when adding/removing nodes.

Nevertheless, there are some types of networks for which a sharp classification of nodes into communities may be elusive, such as when communities overlap (see section 11 in [4]), a situation motivating the development of the method of Ahn *et al*. [32]. To illustrate this point we perform an explicit comparison between the method of Ahn *et al*. and functionInk with a specific example. The open question to compare both methods is how to determine partitions over sets of nodes from partitions over sets of links. A possibility comes from the identification of sets of nodes whose links belong to the same set of link partition(s), since this is the idea behind the development of functionInk. Therefore, we should find similar communities (clusters in the nodes’ partition) if we use the method of Ahn *et al*. to identify a partition of links and i) we then identify communities as those nodes sharing links classified in the same clusters in the partition of links or ii) we find these communities directly with functionInk.

To explore this question, we re-analyze the microbial network shown in Fig. 13 with functionInk considering a single type of link and the communities found at the maximum of the total partition density, together with the partition in links determined with the method of Ahn *et al*., using average linkage as a clustering algorithm in both cases. In Fig. 7 we show the network in which, for clarity, we removed communities with less than three members (although we kept some nodes with a high density of links).

**Figure 7:**
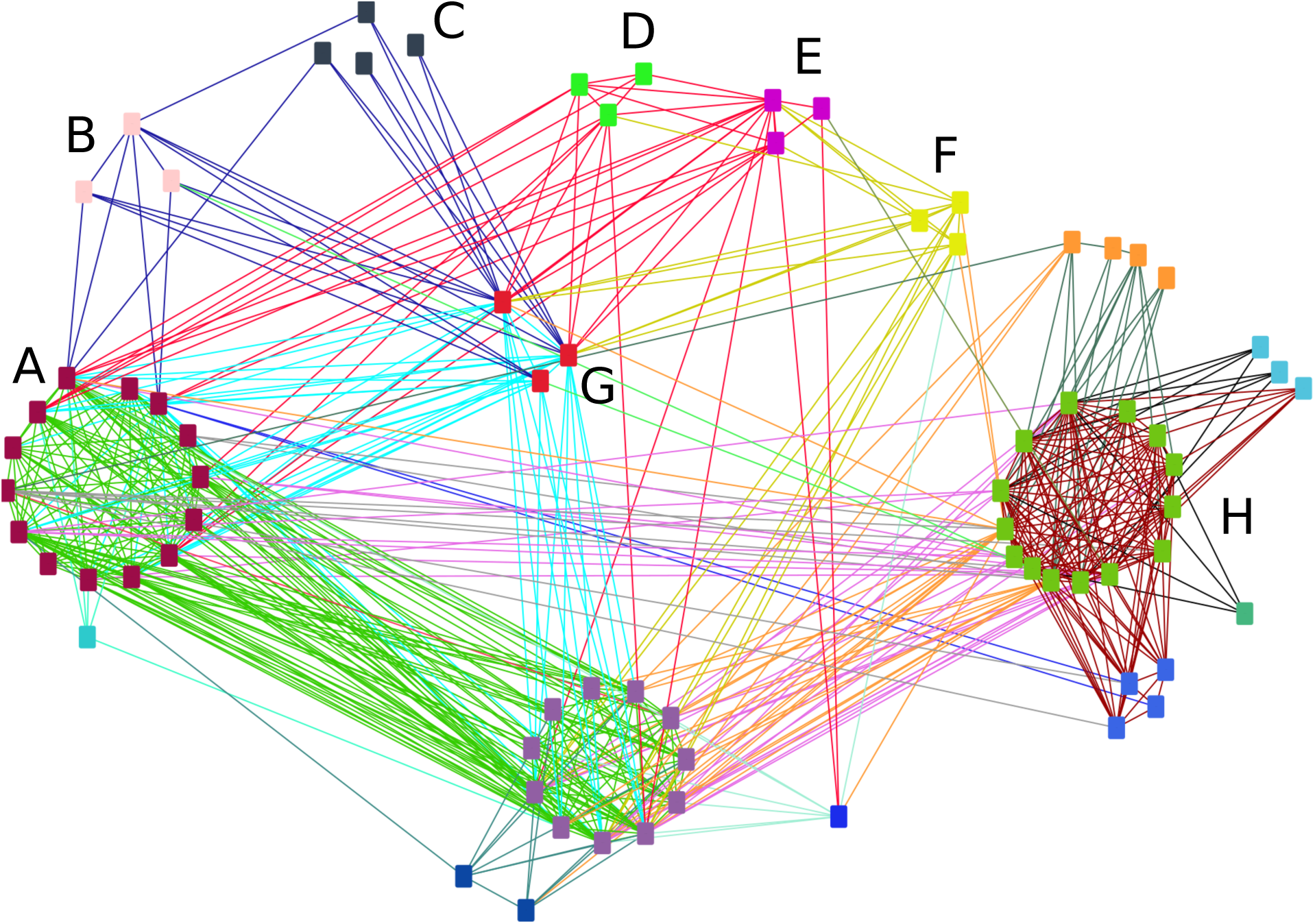
Comparison between functionInk and Ahn *et al*. method. [32]. Microbial network analyzed in Fig. 13 now comparing the partition over the set of nodes found with the total partition density with functionInk (leading to communities of nodes sharing the same color and close in space) and the partition over the set of links found with Ahn *et al*. method (leading to groups of links sharing the same color and connecting similar community nodes). The network is reduced in size and rearranged for the sake of a more clear comparison. Analysis of the communities labeled A-G is found in the text. Note that there is not full correspondence with the communities found in Fig. 13, since we do not use the information from the types of links here.

Partitions defined by Ahn *et al*. method are identified by different colors for the links, and those defined by functionInk by different colors for the nodes (which are further located close in space). It is immediately apparent the good match between both methods, as expected since functionInk departs from the method of Ahn *et al*. For instance, the community labeled G can be defined as the one having cyan, olive, red and blue links. We also observe, however, some differences. Communities B and C are those having only blue links. functionInk splits these nodes in two communities because all three members of community B are linked to the same member in community G, but this is a connection that none of the C-members have, and similarly with respect to two members of community A (although, in this case, one member of C has one such connection). The same applies for communities D and E, which could be defined as those preferentially having red links, but community E is split from D because it has connections with other nodes, in particular with community F. Nevertheless, functionInk is not able to split community H into sub-communities, which is suggested for instance by subsets of nodes having links to several link partitions, e.g. pink and orange links. Even if these communities would be split stopping at an earlier step (for instance at the maximum of the external partition density) thanks to the flexibility provided by the availability of several partition function definitions, it is also clear that functionInk may have difficulties with highly overlapping communities, a situation in which it may be useful to combine both methods.

### Clustering algorithm

After computing the similarity between nodes with the method presented in the Results, the algorithm clusters nodes using one of three hierarchical clustering algorithms: average linkage [48], single linkage and complete linkage. Starting from each node being a separate cluster, at each step *t* all algorithms join the two most similar clusters *A* and *B*, and compute the similarity between the new combined cluster and all other clusters *C* in a way that depends on the clustering algorithm.

Single linkage is the most permissive algorithm, because the similarity it assigns to the new cluster is the maximum similarity between the two clusters joined and clusters *C*:

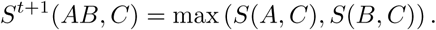

where *t* labels the step of the algorithm, *A* and *B* are the clusters that are joined, *AB* denotes the new composite cluster, and *C* is any other cluster. On the other hand, complete linkage is the most restrictive, assigning the minimum similarity

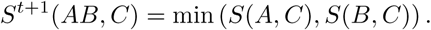

Finally, average linkage assigns an intermediate value computed as the weighted average similarity with the two joined clusters

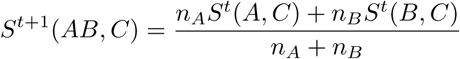

being *n*_*A*_ and *n*_*B*_ the number of elements that *A* and *B* contain, respectively. Identification of the two pools of plants and animals is independent of the clustering method used, but the maximum of the external partition density is achieved earlier for single linkage and later for complete linkage; we found a good compromise between the number and the size of the clusters working with average linkage, in agreement with previously reported results [31], but the clustering method could be selected according to information known from the links. In our experience, single linkage is easily dominated by the giant cluster in high density networks in which modules are prevalent (rather than for guilds). The appropriate clustering method should be guided by the research question. For instance, if gene homology is explored, it is probably more appropriate to use single linkage (as a relative of one gene’s relative is also its relative, i.e. transitivity is automatically fulfilled [36]). On the other hand, if we analyse well-differentiated functional similarity, it might be more appropriate to be conservative and use complete linkage.

### Plant-pollinator networks and topological properties

We selected six plant-pollinator networks artificially generated in [39] with known topological properties, summarized in Table. We consider as topological properties the connectance (fraction of links) of the mutualistic matrix, the connectance of the competition matrices, and the definition of nestedness provided in [40]. Given a mutualistic matrix 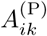 representing presence-absence of interaction between the set of plants, indexed by *i*, and the set of animal species, indexed by *k*, we compute the degree of a species as 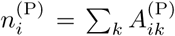 (see Ref. [40] in Supplementary Material). A similar definition would apply for animals 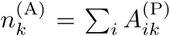. Next we define the ecological overlap between two species of plants *i* and *j* as the number of insects that pollinate both plants:

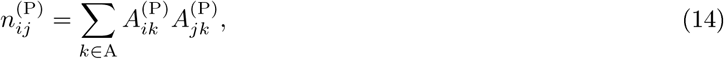

a definition that is equivalent to the numerator Jaccard similarity used in this work. Summing over every pair of plants and normalizing leads to the definition of nestedness:

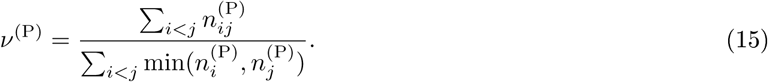

A symmetric definition applies for animals, so we take as final definition of nestedness *ν* = max(*ν*^(P)^, *ν*^(A)^). Note that in Eq. 14, *k* indexes animal species and, since the two pools of plants and animals are separated until the very last steps, changes in the nestedness will have an effect only on the external partiton density.

**Table 1:** Topological properties of the bipartite networks analyzed. Different combinations of nestedness (*ν*), intra-pools connectance *κ*_comp_ and inter-pools connectance *κ*_mut_ were analyzed.

| $\nu$ | $\kappa_{\text{mut}}$ | $\kappa_{\text{comp}}$ |
| --- | --- | --- |
| 0.15 | 0.08 | 0 |
| 0.35 | 0.16 | 0.15 |
| 0.6 | 0.28 | 0.5 |

### Trophic networks

We downloaded the network and metadata provided in [41] and compared the clusters found with those obtained by functionInk. After computing the Tanimoto coefficients as explained above, we cluster the nodes and retrieve the classification found at each step. We then computed five indexes (Rand, Fowlkes and Mallows, Wallace 10, Wallace 01 and Jaccard), implemented in the R PCI function of the PROFDPM package [42]. In order to assign a significance value for the different indexes we obtained, for each index *x*, a bootstrapped distribution with mean 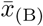 and standard deviation *σ*_(B)_, re-sampling with replacement the samples and recomputing the indexes 10^3^ times. Next we computed 10^3^completely random classifications, obtained by shuffling the identifiers relating each sample with one of the classifications, and retrieving the maximum *x*_(R)_. We finally verified that the random value was significantly different from the bootstrapped distribution by computing the z-scores:

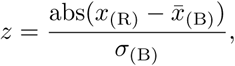

which we considered significant if it was higher than 2.5. Heatmaps were generated with the HEATMAP.2 function in R package GPLOTS.

### Bacterial networks

We considered a public dataset of 753 bacterial communities sampled from rainwater-filled beech tree-holes (Fagus spp.) [44], leading to 2874 Operative Taxonomic Units (OTUs) at the 97% of 16 rRNA sequence similarity. These communities were compared with Jensen-Shannon divergence [45], and automatically clustered following the method proposed in Ref. [49] to identify enterotypes. The clusters found with this method in [43] were used to color the community labels in Fig. 13.

The inference of the OTU network started quantifying correlations between OTUs abundances with SparCC [46]. To perform this computation, from the original OTUs we reduced the data set removing rare taxa with less than 100 reads or occurring in less than 10 samples, leading to 619 OTUs. Then, the significance of the correlations was evaluated bootstrapping the samples 100 times the data and estimating pseudo p-values for each of the *N* (*N* − 1)*/*2 pairs. A relationship between two OTUs was considered significant and represented as a link in the network if the correlation was larger than 0.2 in absolute value and the pseudo p-value lower than 0.01. The network obtained in this way was analyzed using Cytoscape [50].

In Suppl. Fig. 13 we show the network obtained with the partitions found at the maximum of the external, total and internal partition densities. The combination of the three partitions allow us to individuate different key players in the network, and its relation with the definition of structural equivalence we adopted. For instance, only one OTU from the green functional group in the partitioning generated with the total partition density, have an important number of co-occurrences with members of the red functional group. In addition, it is the only one having a significant segregation with respect to a highly abundant member of the blue functional groups. These two highly abundant segregating OTUs are *Pseudomonas putida* (green functional group) and *Serratia fonticola* (blue functional group), both of which were shown to dominate two of the *β*−diversity-classes [43]. In the partitioning found with the external partition density, *Pseudomonas putida* appears as an orphan node and the functional group of *Serratia fonticola* is split in three. This example illustrates how the different stopping criteria can be used to individuate larger or smaller differences in the connectivity.

## Supplementary Figures

**Figure 8:**
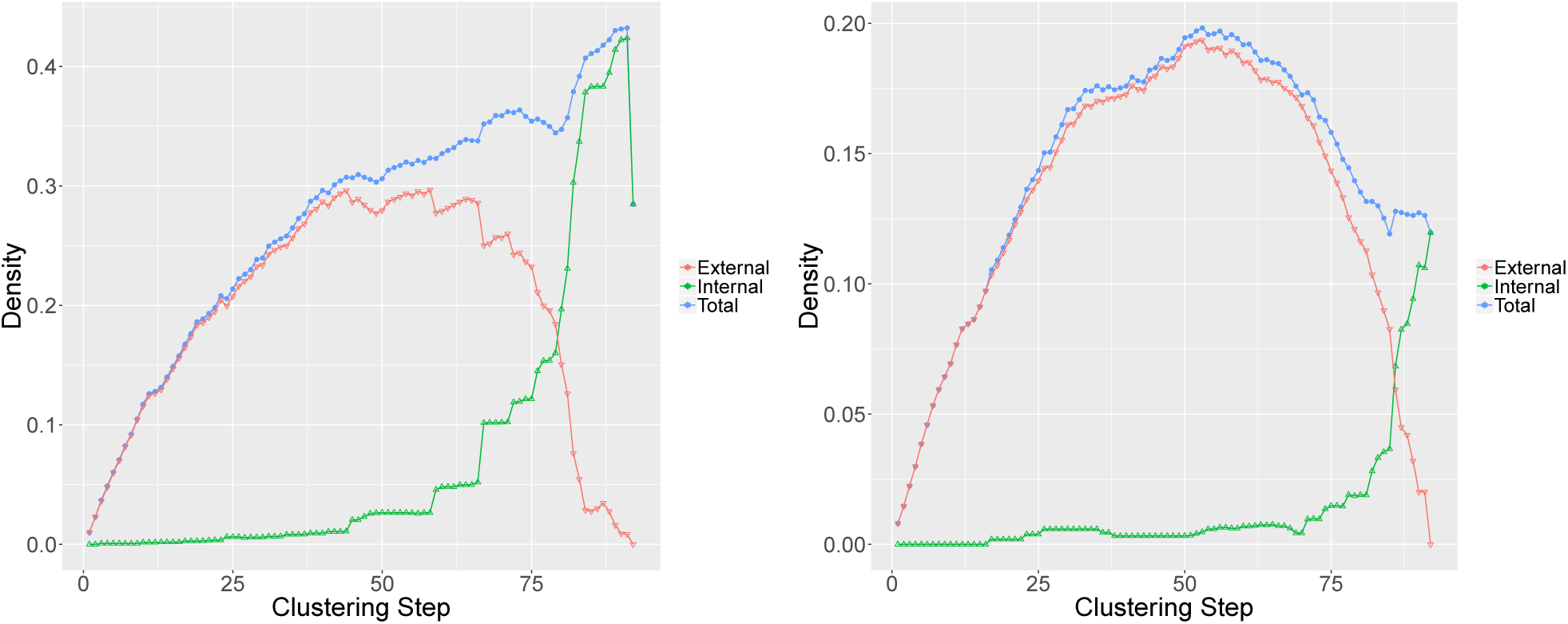
Partition densities of synthetic mutualistic networks. Networks with nestedness *ν* = 0.15, *κ*_mut_ = 0.08, and *κ*_comp_ = 0.5 (left) or *κ*_comp_ = 0.15 (right). Changing the connectance change the relative value between the external and internal partition densities.

**Figure 9:**
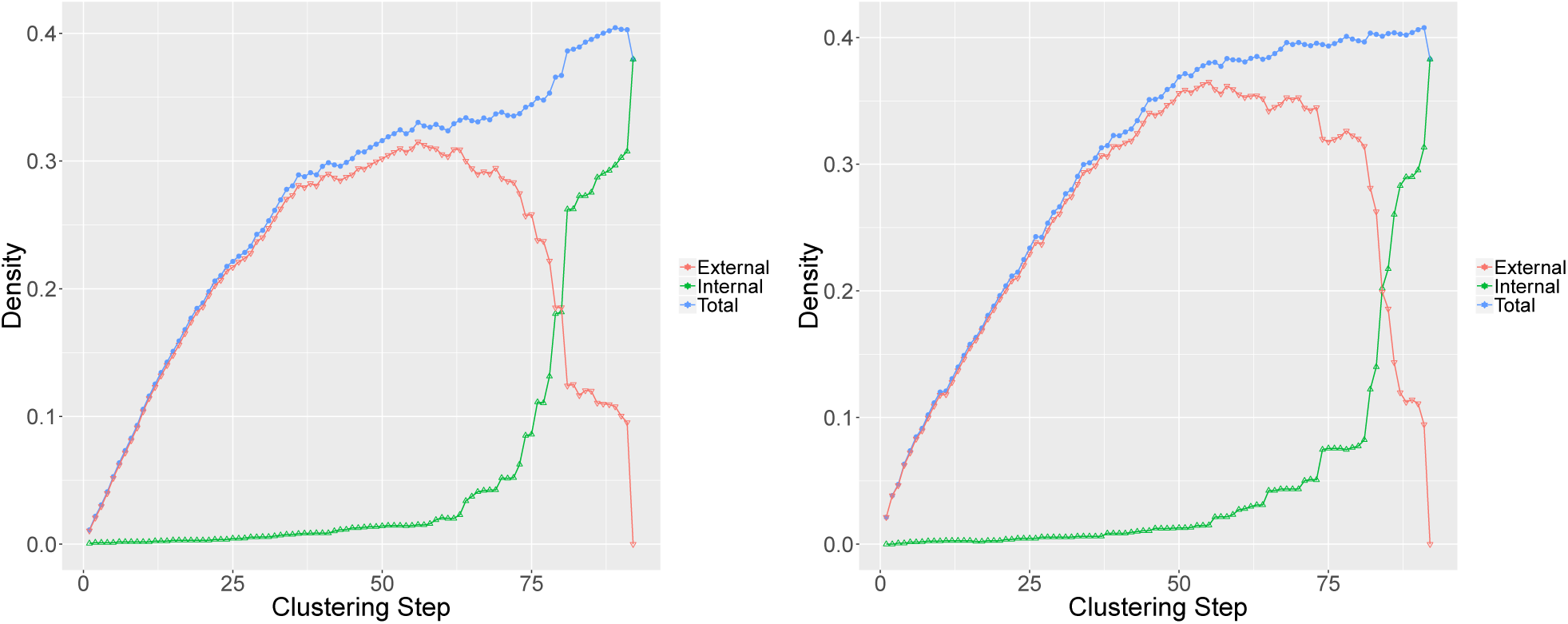
Partition densities for synthetic mutualistic networks. Networks with *κ*_comp_ = 0.5, *κ*_mut_ = 0.28 and *ν* = 0.35 (left) or *ν* = 0.6 (right). The high connectance of both networks make the internal partition density dominant, and two pools are detected through the total partition density. Nevertheless, the increase of the nestedness is detected through an increase in the external partition density, which makes the second network more disassortative.

**Figure 10:**
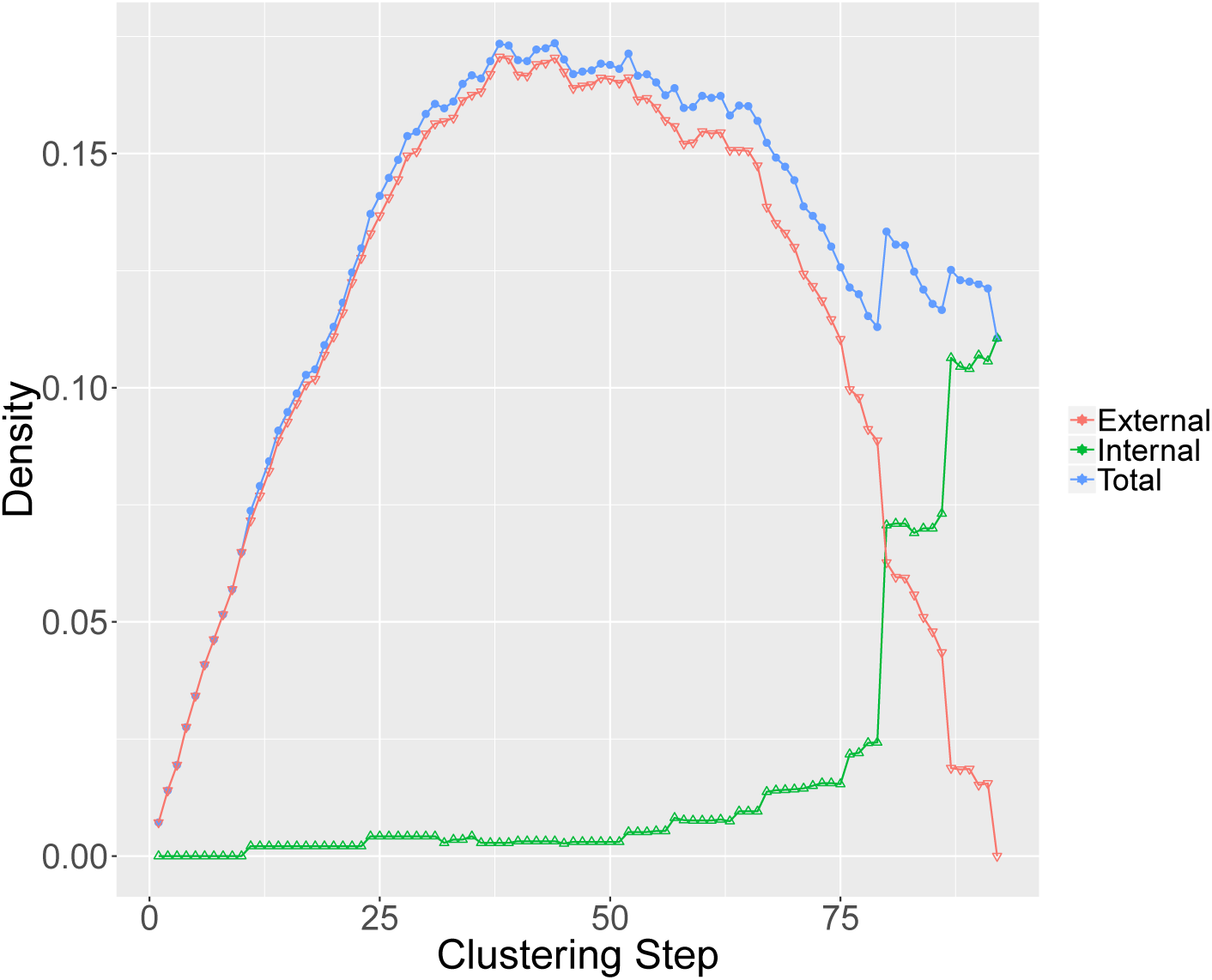
Partition densities of synthetic mutualistic networks. Network with nestedness *ν* = 0.05, *κ*_mut_ = 0.065 and *κ*_comp_ = 0.15. The low connectance hinders the detection of the two pools of plants and pollinators.

**Figure 11:**
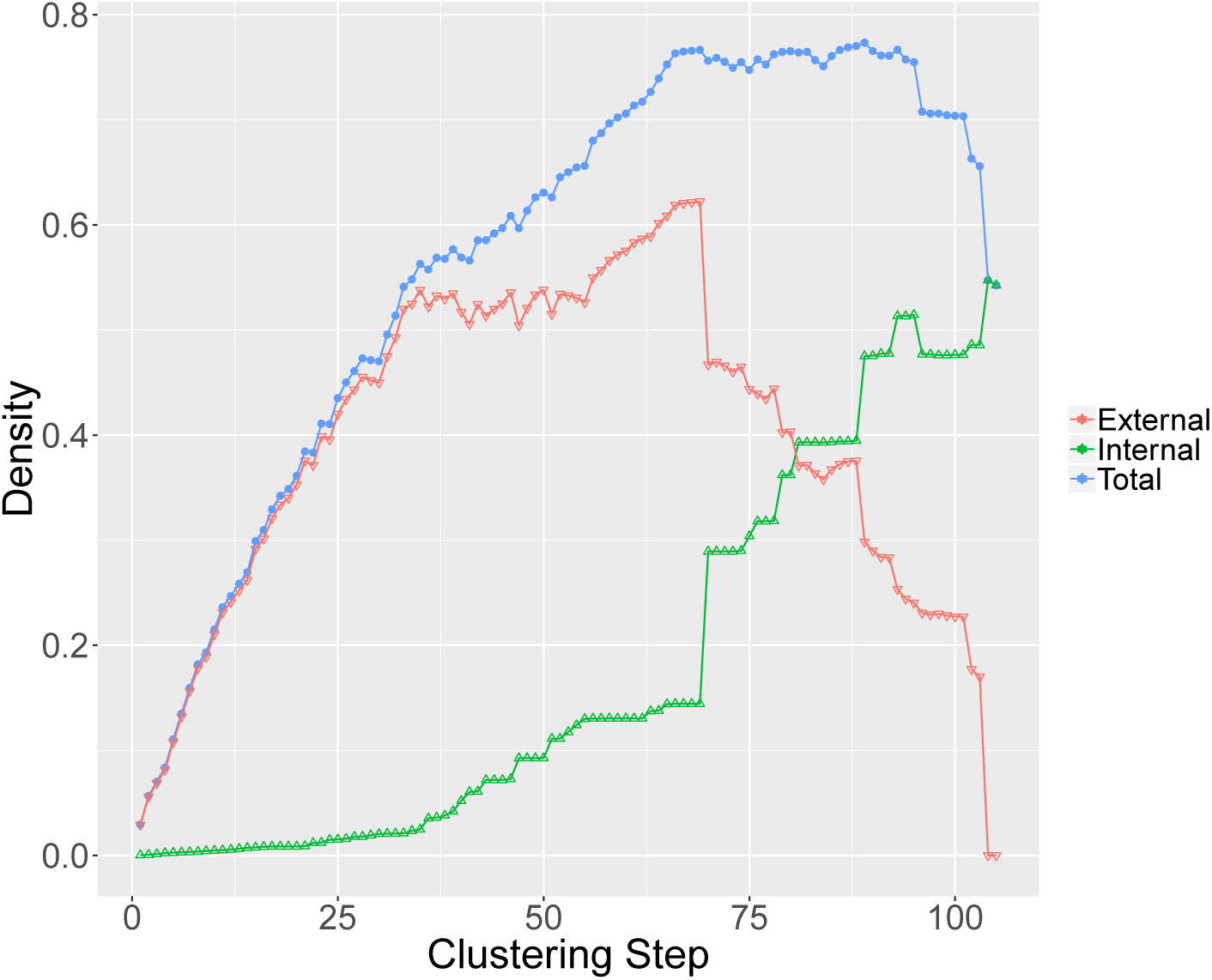
Partition density of the trophic network. The internal partition density peaks when there are three clusters, consistent with the existence of three trophic layers. The external partition density has a maximum at step 69, which is analyzed in detail with respect to the reference classification found in [41].

**Figure 12:**
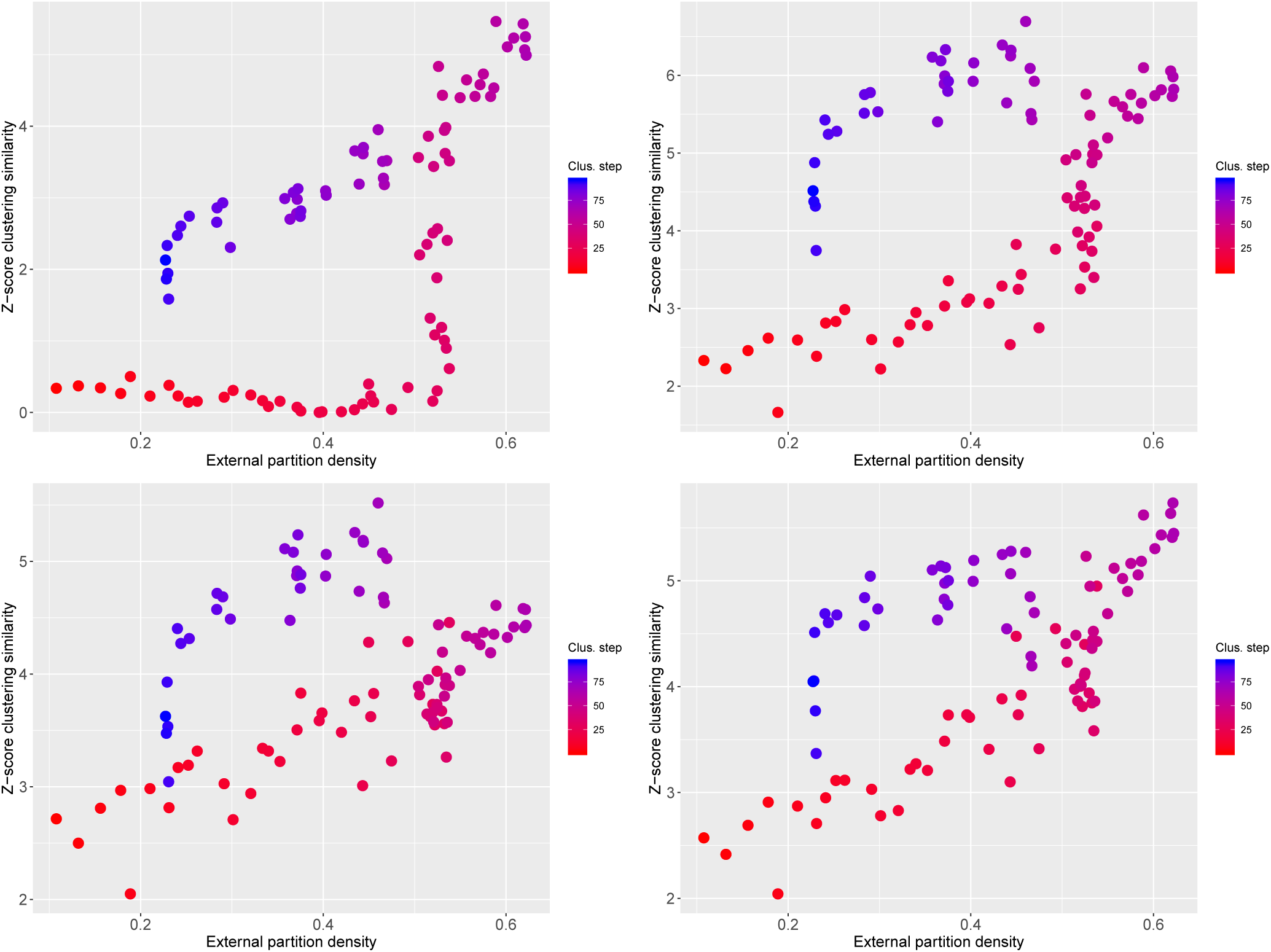
Comparison between classifications of the trophic network. Similarity between the reference classification found in [41] and the one found with functionInk is performed with the Z-score of a different indexes: Wallace 01 (Top left), Fowlkes and Mallows (Top right), Jaccard (Bottom left) and Rand (Bottom right). All indexes bring significant values and the maximum similarity is close to the maximum of the external partition density.

**Figure 13:**
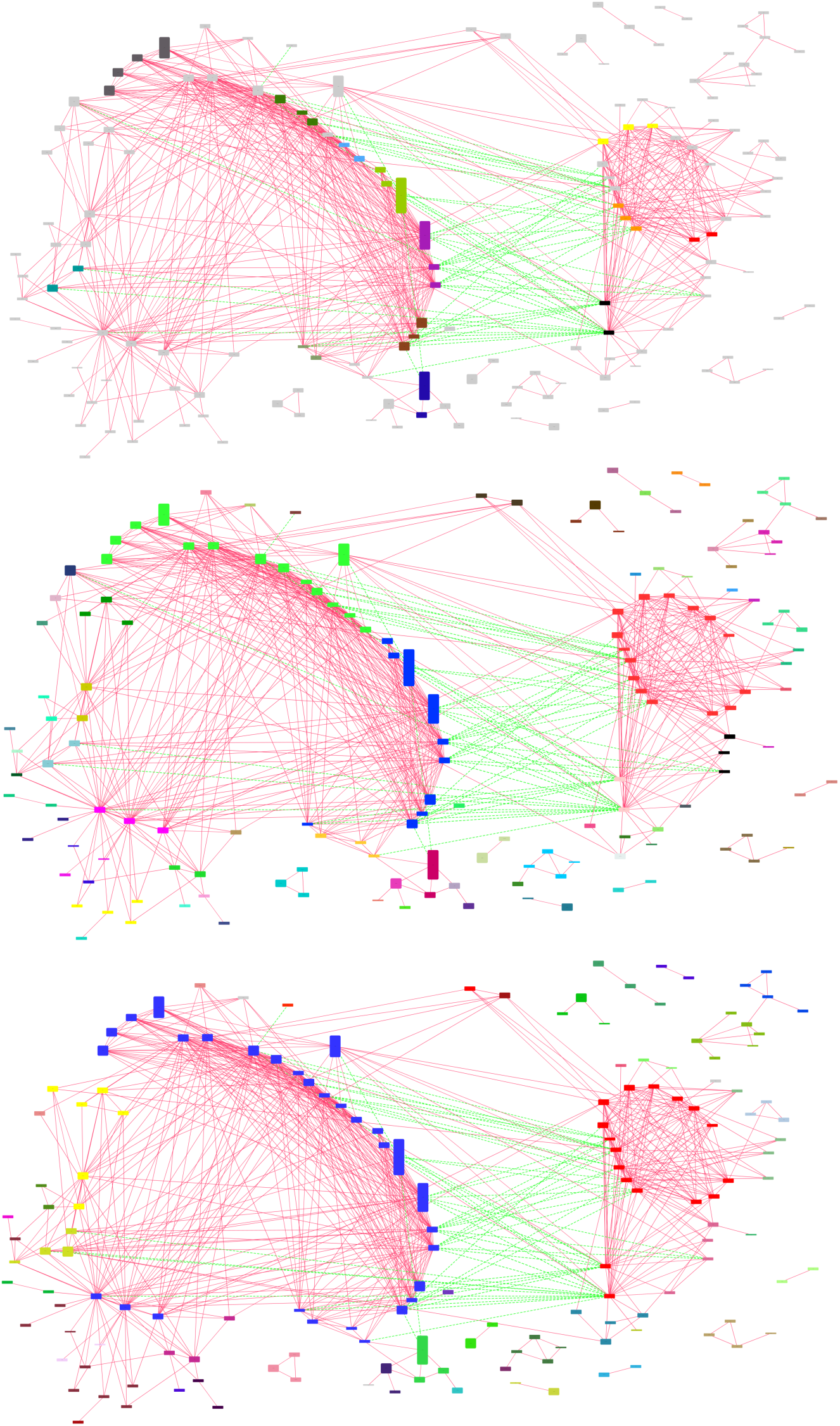
Comparison of functional groups in the microbial network. Network of significant co-occurrences (continuous links) and segregations (dotted links) at the species level (nodes). Colors indicate functional group membership, which was determined by the maximum of the external (top), total (middle) and internal partition densities (bottom). Orphan nodes are colored gray in the top figure for clarity. The higher value of the internal partition density (see Suppl. Fig. 14) suggests that a modular structure is the more appropriate to describe the functional groups. This is confirmed by the low number of guilds (top figure) and the good agreement between the global topological structure and the modules (bottom figure). Communities were automatically located close in space according to the partition found with the total partition density (middle) and blue and green communities rearranged manually to make more clear their connections, in particular we separated one node in the green community being the only one with co-occurrences with other communities on the right-hand side.

**Figure 14:**
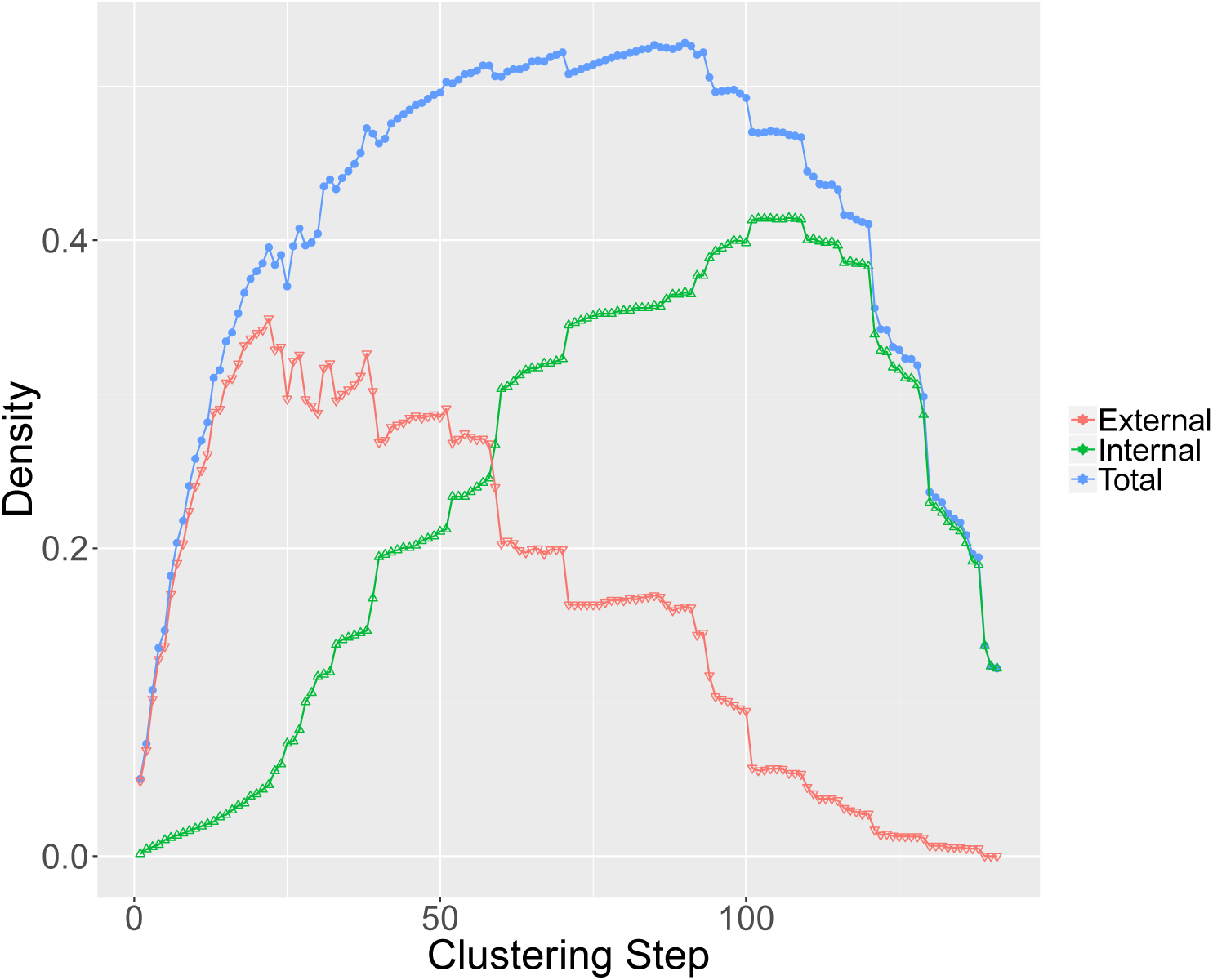
Partition density of the microbial network. The external partition density brings a poor reduction in the complexity of the network, with only 22 elements joined, while the internal partition density achieves a higher value and still a good number of clusters. Results suggest that modules are more relevant in this network given the high number of intra-cluster co-occurrences, later confirmed by visual inspection in the Main Text.

## References

[1] J. E. Cohen and D. W. Stephens, Food webs and niche space. No. 11, Princeton University Press, 1978.

[2] R. MacArthur, “Fluctuations of animal populations and a measure of community stability,” Ecology, vol. 36, p. 533, July 1955.

[3] M. Kivelä, A. Arenas, M. Barthelemy, J. P. Gleeson, Y. Moreno, and M. A. Porter, “Multilayer networks,” Journal of Complex Networks, vol. 2, no. 3, pp. 203–271, 2014.

[4] S. Fortunato, “Community detection in graphs,” Physics Reports, vol. 486, no. 3–5, pp. 75–174, 2010.

[5] S. Allesina and M. Pascual, “Food web models: a plea for groups,” Ecology Letters, vol. 12, no. 7, pp. 652–662, 2009.

[6] S. Pilosof, M. A. Porter, M. Pascual, and S. Kéfi, “The multilayer nature of ecological networks,” Nature Ecology & Evolution, vol. 1, no. 4, p. 0101, 2017.

[7] S. Boccaletti, V. Latora, Y. Moreno, M. Chavez, and D.-U. Hwang, “Complex networks: Structure and dynamics,” Physics reports, vol. 424, no. 4, pp. 175–308, 2006.

[8] A. Carr, C. Diener, N. S. Baliga, and S. M. Gibbons, “Use and abuse of correlation analyses in microbial ecology,” The ISME journal, p. 1, 2019.

[9] A. Pascual-García, J. Tamames, and U. Bastolla, “Bacteria dialog with santa rosalia: Are aggregations of cosmopolitan bacteria mainly explained by habitat filtering or by ecological interactions?,” BMC Microbiology, vol. 14, no. 1, p. 284, 2014.

[10] C. S. Elton, Animal ecology. New York: The Macmillan Company, 1927.

[11] D. Simberloff and T. Dayan, “The guild concept and the structure of ecological communities,” Annual Review of Ecology and Systematics, vol. 22, no. 1, pp. 115–143, 1991.

[12] M. E. Newman and E. A. Leicht, “Mixture models and exploratory analysis in networks,” Proceedings of the National Academy of Sciences, vol. 104, no. 23, pp. 9564–9569, 2007.

[13] M. E. Newman, “Finding community structure in networks using the eigenvectors of matrices,” Physical Review E, vol. 74, no. 3, p. 036104, 2006.

[14] E. Estrada and J. A. Rodríguez-Velázquez, “Spectral measures of bipartivity in complex networks,” Physical Review E, vol. 72, no. 4, p. 046105, 2005.

[15] M. T. Schaub, J.-C. Delvenne, M. Rosvall, and R. Lambiotte, “The many facets of community detection in complex networks,” Applied Network Science, vol. 2, no. 1, p. 4, 2017.

[16] L. Peel, D. B. Larremore, and A. Clauset, “The ground truth about metadata and community detection in networks,” Science advances, vol. 3, no. 5, p. e1602548, 2017.

[17] M. Girvan and M. E. Newman, “Community structure in social and biological networks,” Proceedings of the National Academy of Sciences, vol. 99, no. 12, pp. 7821–7826, 2002.

[18] R. Lambiotte, J.-C. Delvenne, and M. Barahona, “Laplacian dynamics and multiscale modular structure in networks,” arXiv preprint 0812.1770, 2008.

[19] P. J. Mucha, T. Richardson, K. Macon, M. A. Porter, and J.-P. Onnela, “Community structure in time-dependent, multiscale, and multiplex networks,” Science, vol. 328, no. 5980, pp. 876–878, 2010.

[20] M. De Domenico, A. Lancichinetti, A. Arenas, and M. Rosvall, “Identifying modular flows on multilayer networks reveals highly overlapping organization in interconnected systems,” Physical Review X, vol. 5, no. 1, p. 011027, 2015.

[21] M. Bazzi, M. A. Porter, S. Williams, M. McDonald, D. J. Fenn, and S. D. Howison, “Community detection in temporal multilayer networks, with an application to correlation networks,” Multiscale Modeling & Simulation, vol. 14, no. 1, pp. 1–41, 2016.

[22] S. Wasserman and K. Faust, Social network analysis: Methods and applications, vol. 8. Cambridge university press, 1994.

[23] P. W. Holland, K. B. Laskey, and S. Leinhardt, “Stochastic blockmodels: First steps,” Social networks, vol. 5, no. 2, pp. 109–137, 1983.

[24] C. De Bacco, E. A. Power, D. B. Larremore, and C. Moore, “Community detection, link prediction, and layer interdependence in multilayer networks,” Physical Review E, vol. 95, no. 4, p. 042317, 2017.

[25] M. E. Newman, “Equivalence between modularity optimization and maximum likelihood methods for community detection,” Physical Review E, vol. 94, no. 5, p. 052315, 2016.

[26] B. Karrer and M. E. Newman, “Stochastic blockmodels and community structure in networks,” Physical Review E, vol. 83, no. 1, p. 016107, 2011.

[27] T. Valles-Catala, F. A. Massucci, R. Guimera, and M. Sales-Pardo, “Multilayer stochastic block models reveal the multilayer structure of complex networks,” Physical Review X, vol. 6, no. 1, p. 011036, 2016.

[28] M. Ganji, J. Chan, P. J. Stuckey, J. Bailey, C. Leckie, K. Ramamohanarao, and I. Davidson, “Image constrained blockmodelling: a constraint programming approach,” in Proceedings of the 2018 SIAM International Conference on Data Mining, pp. 19–27, SIAM, 2018.

[29] S. P. Borgatti, M. G. Everett, and P. R. Shirey, “LS sets, lambda sets and other cohesive subsets,” Social networks, vol. 12, no. 4, pp. 337–357, 1990.

[30] J. J. Luczkovich, S. P. Borgatti, J. C. Johnson, and M. G. Everett, “Defining and measuring trophic role similarity in food webs using regular equivalence,” Journal of Theoretical Biology, vol. 220, no. 3, pp. 303–321, 2003.

[31] P. Yodzis and K. O. Winemiller, “In search of operational trophospecies in a tropical aquatic food web,” Oikos, pp. 327–340, 1999.

[32] Y.-Y. Ahn, J. P. Bagrow, and S. Lehmann, “Link communities reveal multiscale complexity in networks,” Nature, vol. 466, no. 7307, pp. 761–764, 2010.

[33] R. Guimera and L. A. N. Amaral, “Cartography of complex networks: modules and universal roles,” Journal of Statistical Mechanics: Theory and Experiment, vol. 2005, no. 02, p. P02001, 2005.

[34] S. P. Borgatti and M. G. Everett, “The class of all regular equivalences: Algebraic structure and computation,” Social networks, vol. 11, no. 1, pp. 65–88, 1989.

[35] T. T. Tanimoto, “elementary mathematical theory of classification and prediction,” 1958.

[36] A. Pascual-García, D. Abia, Á. R. Ortiz, and U. Bastolla, “Cross-over between discrete and continuous protein structure space: insights into automatic classification and networks of protein structures,” PLoS Computational Biology, vol. 5, no. 3, p. e100033.1, 2009.

[37] A. Harrer and A. Schmidt, “Blockmodelling and role analysis in multi-relational networks,” Social Network Analysis and Mining, vol. 3, no. 3, pp. 701–719, 2013.

[38] M. G. Everett and S. P. Borgatti, “Exact colorations of graphs and digraphs,” Social networks, vol. 18, no. 4, pp. 319–331, 1996.

[39] A. Pascual-García and U. Bastolla, “Mutualism supports biodiversity when the direct competition is weak,” Nature Communications, vol. 8, p. 14326, Feb. 2017.

[40] U. Bastolla, M. A. Fortuna, A. Pascual-García, A. Ferrera, B. Luque, and J. Bascompte, “The architecture of mutualistic networks minimizes competition and increases biodiversity,” Nature, vol. 458, pp. 1018–1020, Apr. 2009.

[41] S. Kéfi, V. Miele, E. A. Wieters, S. A. Navarrete, and E. L. Berlow, “How structured is the entangled bank? the surprisingly simple organization of multiplex ecological networks leads to increased persistence and resilience,” PLoS Biology, vol. 14, no. 8, p. e1002527, 2016.

[42] M. S. Shotwell et al., “profdpm: An r package for map estimation in a class of conjugate product partition models,” J Stat Softw, vol. 53, no. 8, pp. 1–18, 2013.

[43] A. Pascual-García and T. Bell, “Community-level signatures of ecological succession in natural bacterial communities,” bioRxiv, p. 636233, 2019.

[44] D. W. Rivett and T. Bell, “Abundance determines the functional role of bacterial phylotypes in complex communities,” Nature microbiology, p. 1, 2018.

[45] D. M. Endres and J. E. Schindelin, “A new metric for probability distributions,” IEEE Transactions on Information Theory, vol. 49, pp. 1858–1860, July 2003.

[46] J. Friedman and E. J. Alm, “Inferring correlation networks from genomic survey data,” PLoS Computational Biology, vol. 8, no. 9, p. e1002687, 2012.

[47] M. T. Agler, J. Ruhe, S. Kroll, C. Morhenn, S.-T. Kim, D. Weigel, and E. M. Kemen, “Microbial hub taxa link host and abiotic factors to plant microbiome variation,” PLoS Biology, vol. 14, no. 1, p. e1002352, 2016.

[48] R. R. Sokal, “A statistical method for evaluating systematic relationships,” Univ Kans Sci Bull, vol. 38, pp. 1409–1438, 1958.

[49] M. Arumugam, J. Raes, E. Pelletier, D. Le Paslier, T. Yamada, D. R. Mende, G. R. Fernandes, J. Tap, T. Bruls, J.-M. Batto, et al., “Enterotypes of the human gut microbiome,” Nature, vol. 473, no. 7346, pp. 174–180, 2011.

[50] P. Shannon, A. Markiel, O. Ozier, N. S. Baliga, J. T. Wang, D. Ramage, N. Amin, B. Schwikowski, and T. Ideker, “Cytoscape: a software environment for integrated models of biomolecular interaction networks,” Genome research, vol. 13, no. 11, pp. 2498–2504, 2003.

